# Gene expression changes in *Drosophila melanogaster* females associated with the rescue of the *bag of marbles* (*bam*) hypomorph fertility defect by *Wolbachia pipientis*

**DOI:** 10.1101/2023.09.22.558898

**Authors:** Catherine H. Kagemann, Gabriela M. Colocho, Charles F. Aquadro

## Abstract

*Wolbachia pipientis* are maternally transmitted endosymbiotic bacteria commonly found in arthropods and nematodes. These bacteria manipulate reproduction of the host to increase their transmission using mechanisms, such as cytoplasmic incompatibility, that favor infected female offspring. Interestingly, *W. pipientis* infection partially rescues female fertility in flies containing hypomorph mutations of *bag of marbles* (*bam*) in *D. melanogaster*, which functions in germline stem cell (GSC) maintenance and differentiation. Here we evaluated the effect of *W. pipientis* variant infection on host gene expression using RNA-seq. RNA-seq analysis using the *W. pipientis* infected relative to uninfected *bam* hypomorph genotype had differential expression for many of *bam’s* genetic and physical interactors. The highest magnitude of differential expression of *bam’s* interactors and other GSC genes involved in cystoblast differentiation occurred prior to peak fertility rescue. Our analysis showed that ubiquitination and histone lysine methylation could be important mechanisms involved in the *bam* hypomorph fertility rescue by *W. pipientis*.

## Introduction

In female *Drosophila melanogaster*, the *bag of marbles* (*bam)* gene product is essential for germline stem cell (GSC) renewal and cystoblast differentiation [1–3]. The *bam* hypomorph single amino acid protein coding mutant *D. melanogaster* contain tumorous ovaries due to over proliferation of GSC-like cells, resulting in partial sterility in females [1, 4]. A maternally inherited bacterial endosymbiont, *Wolbachia pipientis* infects many insects and manipulates host reproduction to increase its transmission, using mechanisms such as cytoplasmic incompatibility, male killing, and feminization (3–6). Interestingly, *W. pipientis* infection of female *bam* hypomorph flies causes a partial rescue of the otherwise reduced hypomorph fertility and a reduction in over-proliferating GSC-like cells. In contrast to females, the phenotype of male *bam* hypomorphs are fully sterile and this phenotype is not rescued by infection by *W. pipientis* [5].

As *W. pipientis* titer is known to influence a wide array of host phenotypes [4, 6, 7], we hypothesized that *W. pipientis* titer could impact *bam* hypomorph fertility rescue. Genetically diverse variants of *W. pipientis* have been characterized in *D. melanogaster* and fall into two categories: *w*Mel-like variants (clades III, VIII) and *w*MelCS-like variants (clade IV) [7, 8]. In *D. melanogaster* males, *w*Mel-like *W. pipientis* variants are associated with a longer host lifespan and low *W. pipientis* titer, while *w*MelCS-like variants are associated with a shorter host lifespan and a higher *W. pipientis* titer in comparison [7]. As *D. melanogaster* males age, *w*MelCS-like *W. pipientis* titer increases while *w*Mel-like titer remains relatively constant [9]. The characterization of *w*Mel-like and *w*MelCS-like *W. pipientis* titer and fly lifespan characteristics have not been conducted in female flies.

We evaluated whether *W. pipientis* titer increases in females as it does in males as the host ages. Next, we identified whether changes in *W. pipientis* titer correlated with changes in the *bam* hypomorph fertility rescue. Utilizing *W. pipientis* variant titer in aged female *D. melanogaster* to maximize *bam* hypomorph fertility rescue, we identified the genes with the highest expression during the rescue.

Using two *W. pipientis* variants known to have a high titer in male *D. melanogaster* and two with a low titer in males, we tested whether 1) *W. pipientis* titer and host age influence the *bam* hypomorph fertility rescue as a means of maximizing rescue-specific gene enrichment and 2) whether different *W. pipientis* variants cause differential rescue of the *bam* hypomorph phenotype at the transcript level as the female host fly ages.

Given a significant amount of evidence showing that *W. pipientis* modulates host gene expression during infection [9–11], we studied the *bam* hypomorph fertility rescue at the transcript level. We hypothesized that two *W. pipientis* groups (*w*Mel-like and *w*MelCS-like) confers distinct gene expression profiles in the host to rescue the *bam* hypomorph phenotype. When *bam* does not function properly, as in the *bam* hypomorph, *W. pipientis* infection could manipulate *bam* gene expression itself or that of *bam’s* genetic and/or physical interactors during the *bam* hypomorph rescue to promote oogenesis. *W. pipientis* could additionally act upstream or downstream of *bam*, completely bypassing *bam* and its interactors to rescue the *bam* hypomorph fertility phenotype.

## Results

### Increased ovarian W. pipientis titer correlates with peak bam hypomorph fertility rescue in six-day old females

We measured fertility of three-, six-, and nine-day old wild-type (WT) uninfected *D. melanogaster* as well as those infected with either of two *w*Mel-like variants (*w*Mel2a and *w*Mel3) and either of two *w*MelCS-like variants (*w*MelCS2a and *w*MelCS2b). Peak fertility occurred in six-day old WT female flies (Fig 1A and S1 Table). Fertility differed significantly between infected and uninfected WT females in three- and nine-day old flies (Fig 1A, Poisson response distribution, p < 0.05) but not in six-day old flies. Further, *w*Mel-like infected *Drosophila* fertility was statistically different from *w*MelCS-like infected WT *Drosophila* at all ages (Fig 1A, Poisson response distribution, p < 0.05).

**Fig 1.**
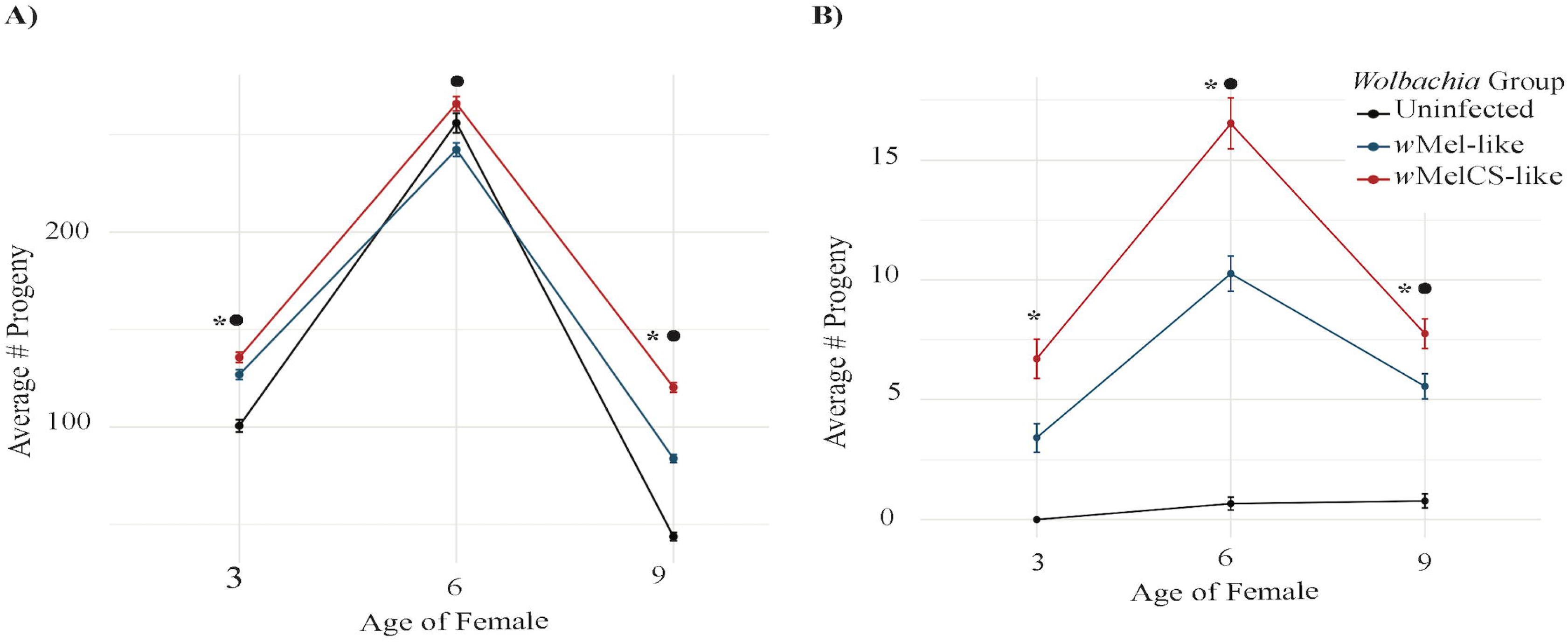
Fertility in the *bam* hypomorph flies shows that fertility rescue is dependent on the age of the female fly and *W. pipientis* genotype. The estimated marginal mean progeny per *W. pipientis* group (*w*Mel-like: *w*Mel2a and *w*Mel3, *w*MelCS-like: *w*MelCS2a and *w*MelCS2b) infecting *D. melanogaster* and the uninfected *D. melanogaster* control in the (A) WT and (B) *bam* hypomorph genotypes. The x-axis shows the age of the female fly after a day of mating, the y-axis (note the difference in y-axes between figures a and b) shows the mean progeny laid, and the bars represent the standard error of each *W. pipientis* group at a given age. The symbol, •, denotes a statistically significant difference between the uninfected control and the *W. pipientis* infected samples at each given age. The symbol, denotes a statistically significant difference between *W. pipientis* groups (*w*Mel-like versus *w*MelCS-like) at each given age.

In contrast to *bam* WT flies, the fertility of *W. pipientis*-infected *bam* hypomorph flies exhibited significant differences compared to uninfected flies in six- and nine-day old flies, suggesting a fertility rescue effect by *W. pipientis* (Fig 1B, Poisson response distribution, p < 0.05). Peak fertility rescue occurred in six-day old *bam* hypomorph flies but never reached *bam* WT fertility levels (Fig 1 and S1 Table). Fertility also differed between *bam* hypomorph females infected with *w*Mel-like and *w*MelCS-like *W. pipientis* group at all ages (Fig 1B, Poisson response distribution, p < 0.05).

Our experimental design allowed us to quantify fertility, titer, and RNA expression from the same female flies. We focused on ovarian titer in females due to our interest in how *W. pipientis* was rescuing female fertility of a *bam* hypomorph. We first assessed titer and fertility in WT flies infected with *w*Mel2a, *w*Mel3, *w*MelCS2a, or *w*MelCS2b *W. pipientis* variants. Combining the data for the four *W. pipientis* variants, we observed a positive relationship between female fertility and ovarian titer in the *bam* WT genotype across all ages (S1 Fig, linear regression model, p > 0.05). None of these correlations were statistically significant (S1 Fig, linear regression model, p > 0.05). Next, we asked whether the rescue of female fertility is impacted by *W. pipientis* titer overall in the *bam* hypomorph ovaries as the flies age. We found that *W. pipientis* titer does impact rescue of six- and nine-day old *bam* hypomorph females (p < 0.05), but not three-day old females (p > 0.05) (Fig 2). Based on the association between the highest fertility rescue and highest titer in six-day-old female flies, we conducted additional studies with the premise that maximum hypomorph *W. pipientis* rescue is occurring at this age.

**Fig 2.**
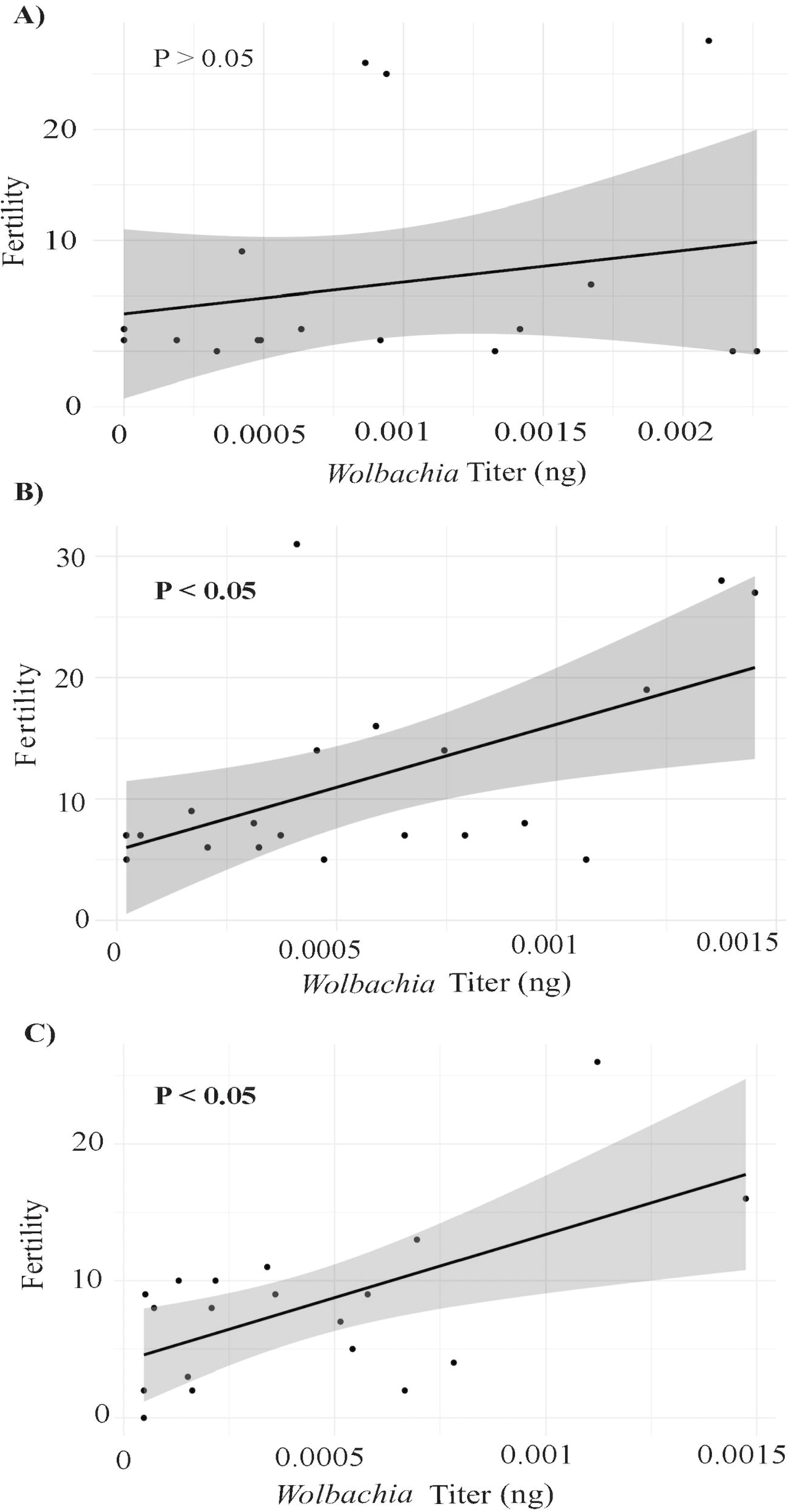
*W. pipientis* titer increases as *bam* hypomorph fertility increases during peak fertility rescue. *Bam* hypomorph fertility and titer of all *W. pipientis* variants infecting *D. melanogaster*. Fertility and titer of mated aged flies: (A) three-day, (B) six-day and (C) nine-day old females. The p-value indicates whether fertility and titer are statistically correlated (linear regression model).

### *W. pipientis* infection alters transcript levels of *bam’s* genetic/physical interactors, but not of *bam* itself in the *bam* hypomorph

We assessed whether *W. pipientis* alters gene expression of *bam* and/or *bam’s* genetic and/or physical interactors to compensate for *bam’s* partial loss of function in the hypomorph by conducting RNA-seq of ovaries from the parent female flies used for *D. melanogaster* fertility and *W. pipientis* titer assays.

Differential expression in *W. pipientis* infected WT flies compared to an uninfected control shows that *W. pipientis* infection alters expression of *bam* and *bam’s* genetic and physical interactors in solely mated six-day old flies (Fig 3A and S2 Table, inferred using DESeq2, p < 0.05). In contrast, in the *bam* hypomorph we observed differential expression for some, but not all, of *bam*’s known genetic and physical interactors between the *W. pipientis* infected and uninfected females in unmated three-, mated three-, and mated six-day old flies (inferred using DESeq2, p < 0.05, Fig 3B). *Bam* itself is only differentially expressed in unmated three-day old *bam* hypomorph flies infected with *w*MelCS2a (S2 Table, [12].)

**Fig 3.**
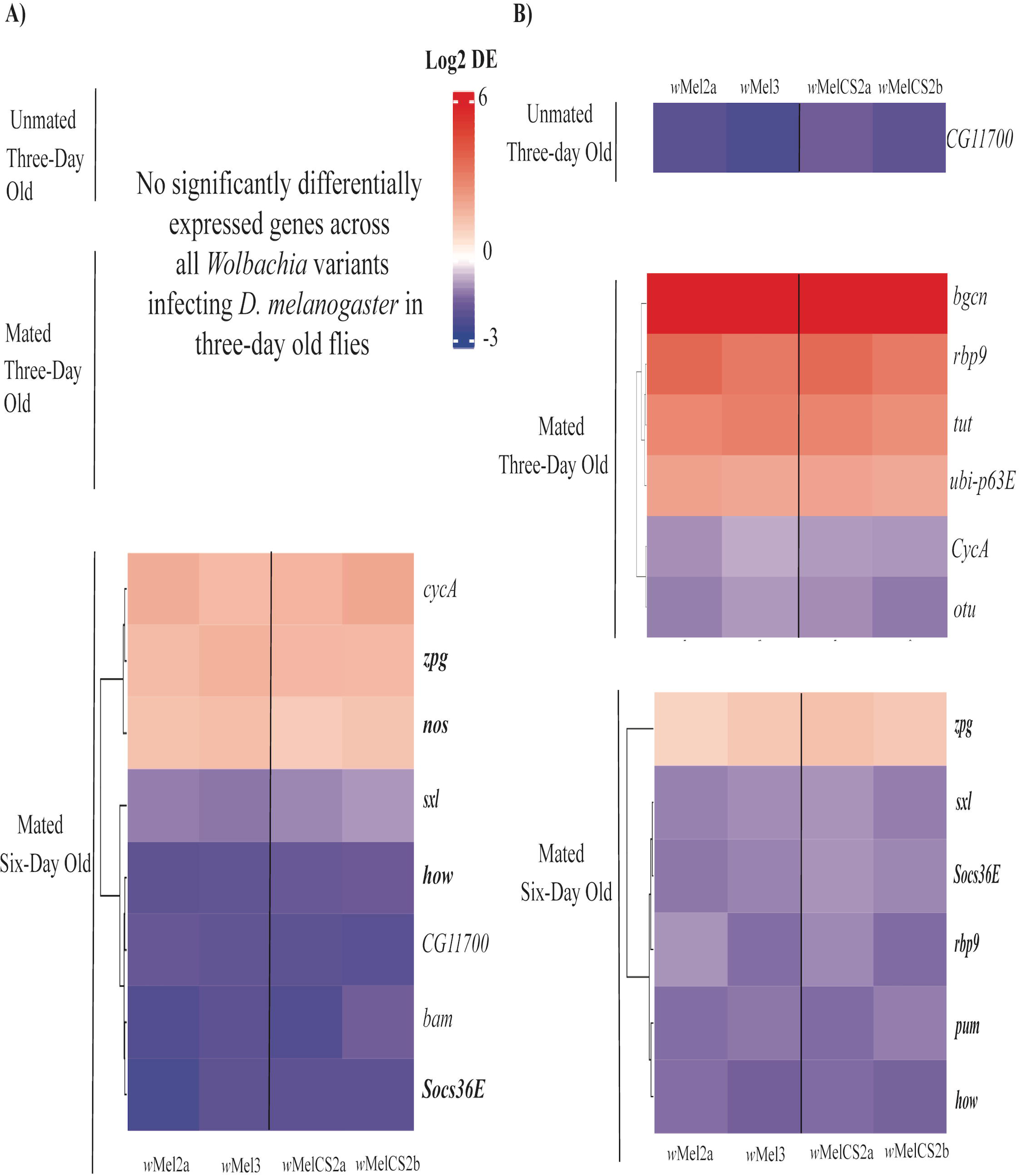
*W. pipientis* infection causes differential expression of *bam’s* genetic and physical interactors in the *bam* hypomorph. Differential expression of *bam* and *bam’s* genetic/physical interactors in *W. pipientis* infected *D. melanogaster* compared to uninfected *D. melanogaster* in the A) WT and B) *bam* hypomorph genotypes at different age/mating statuses (p < 0.05, abs log2fold change > 1). Each column represents the *W. pipientis* variant infecting the *D. melanogaster* shown at the bottom with a line separating the *w*Mel-like and *w*MelCS-like variants. Genes in bold represent that they are differentially expressed in both genotypes. Only genes differentially expressed across all *W. pipientis* infecting the WT or *bam* hypomorph genotype are shown (refer to S2 Table for all differentially expressed genes).

Within genotypes, there are subtle differences in tissue size, particularly in the *bam* hypomorph genotype due to *W. pipientis* rescuing fertility at a greater magnitude in the *w*MelCS-like compared to the *w*Mel-like variants infecting *D. melanogaster*. Therefore, genes expressed due to differences in tissue size were identified computationally along with genes impacted by age and mating. Genes predicted to be influenced by tissue size, age, and mating in the WT and *bam* hypomorph genotypes are listed in the supplement (S3 Table); they were not removed from our analyses as *W. pipientis* infection might also influence them. Of *bam’s* 36 known genetic and physical interactors, two, *cycA* and *dl*, are differentially expressed and predicted to be impacted by tissue size in mated three-day old *bam* hypomorph flies and none in six-day old *bam* hypomorph flies (p < 0.05, absolute log2fold change > 1, S2 Table and S3 Table). None of *bam’s* genetic and physical interactors were impacted by tissue size in mated three- and six-day old WT flies (p < 0.05, absolute log2fold change > 1, S3 Table). None of *bam’s* genetic or physical interactors are influenced by age or mating in the *bam* hypomorph or WT genotypes according to our predictions (S2 Table and S3 Table). There are additional *bam* interactors that were differentially expressed and predicted to be influenced by tissue size, age and mating when removing the log2fold change cutoff (S2 Table and S3 Table). We hypothesized that genes with a significant effect size are more likely to play a role in the fertility rescue of the *bam* hypomorph. Therefore, our analysis mainly targeted genes exhibiting an absolute log2fold change greater than 1 (S2 Table and S3 Table).

### *Bam* hypomorph-specific differential expression of genes involved in GSC daughter cell differentiation into cystoblast is observed prior to the peak of fertility rescue

To understand whether the *bam* hypomorph fertility rescue by *W. pipientis* correlated with the differential expression of genes other than *bam’s* direct physical and genetic interactors, we determined whether 366 genes known to be involved in GSC differentiation, maintenance, or other processes in oogenesis [13] are differentially expressed in our WT and *bam* hypomorph datasets (S2 Fig). We created gene networks of the differentially expressed genes from Yan et al. (2014) and *bam’s* differentially expressed interactors (Fig 4).

**Fig 4.**
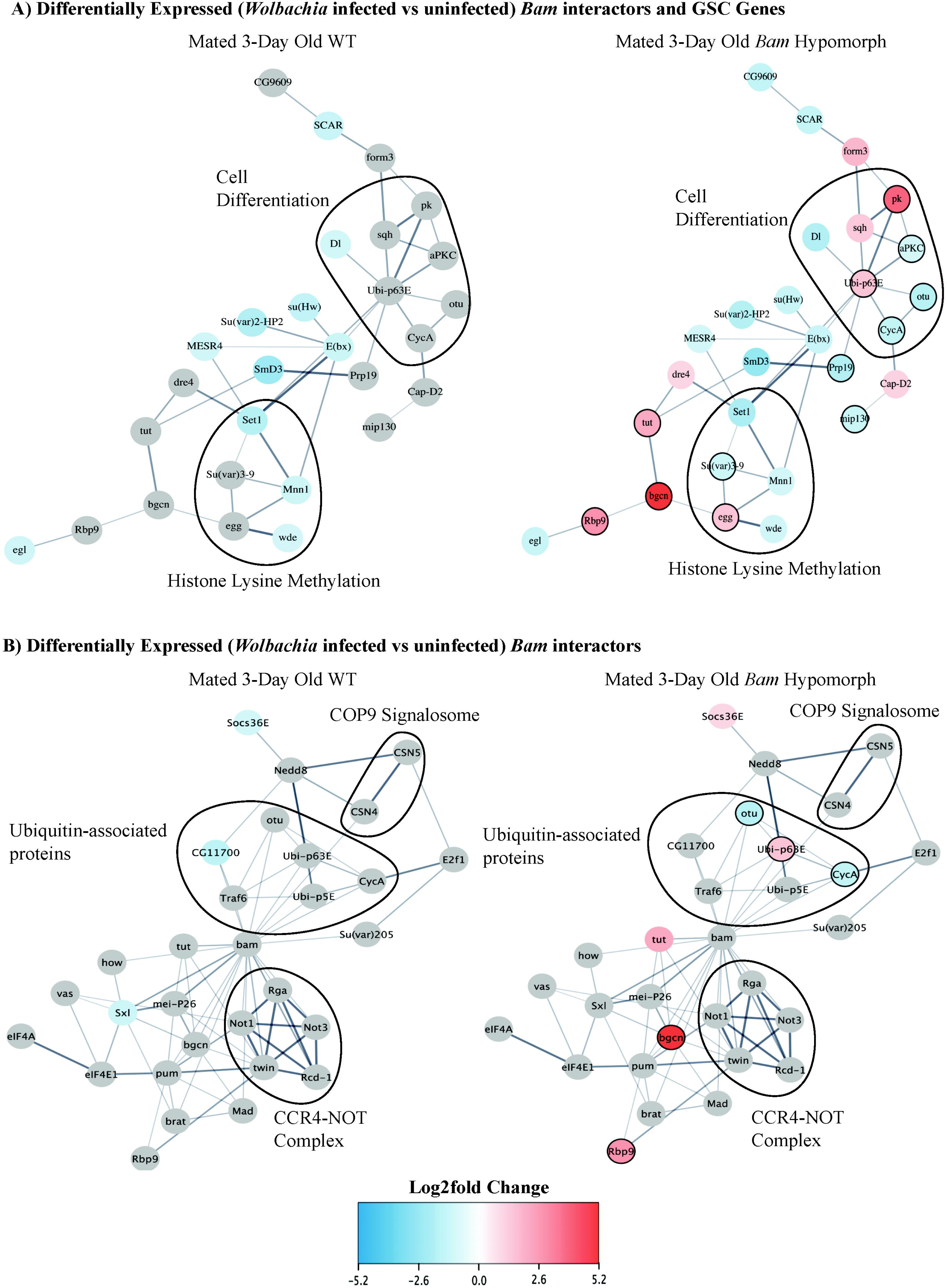
Differential expression of *bam’s* interactors and GSC genes shows genes involved in cystoblast differentiation, histone lysine methylation, and ubiquitination. A) *Bam’s* differentially expressed physical interactors and GSC genes identified from Yan et al. (2014) in mated three-day old WT and *bam* hypomorph flies. The average log2fold change of each differentially expressed gene between flies infected with each *W. pipientis* variant was used as there was not a significant difference between differential expression of these genes in *Drosophila* infected with *w*Mel-like and *w*MelCS-like variants. Genes that are only differentially expressed in the *bam* hypomorph are circled with a black border. The top GO categories were circled within each network. B) Solely Bam’s physical interactors in the mated three-day old WT and *bam* hypomorph genotypes. Three of the largest protein complexes known to be associated with Bam are circled, COP9 Signalosome, Ubiquitin-associated proteins, and the CCR4-NOT complex. Bam’s interactors that are not differentially expressed are annotated with grey circles.

We were particularly interested in the differentially expressed genes specific to the *bam* hypomorph as we hypothesized that they could correlate with the *bam* hypomorph fertility rescue by *W. pipientis*. Interestingly, out of the differentially expressed genes specific to the *bam* hypomorph, the greatest number of genes that are involved in GSC daughter cell differentiation into cystoblast occurred in mated three-day old flies, prior to peak fertility rescue (Fig 4A). Of the *bam* hypomorph-specific differentially expressed genes in three-day old flies, six genes fell into the cell differentiation GO category and four are involved in histone lysine methylation, a process known to be associated with GSC daughter cell differentiation into cystoblast (Fig4A, [14–17].

Of the differentially expressed GSC genes, zero were predicted to be impacted by tissue size in mated three-day old WT or *bam* hypomorph flies (p < 0.05, absolute log2fold change > 1, S2 Table, S3 Table). No GSC genes were influenced by age or mating in the WT genotype and one (*glu*) in the *bam* hypomorph genotype according to our predictions (p value < 0.05, absolute log2fold change > 1, S2 Table and S3 Table).

A separate gene network solely including *bam’s* physical interactors showed that three out of six differentially expressed genes were involved in ubiquitination or deubiquitination in unmated three-day old *bam* hypomorph flies but not WT flies (Fig 4B). No differentially expressed genes were identified among Bam’s physical interactors that are known to be involved in the CCR4-NOT complex or the COP9 signalosome, both crucial for germ stem cell (GSC) self-renewal (Fig 4B).

### Gene enrichment of genes involved in oogenesis, reproduction, and oocyte development/differentiation occurs prior to peak fertility rescue in the *bam* hypomorph

To determine if there were specific classes of genes enriched in the *bam* hypomorph that could be correlated with the *bam* hypomorph fertility rescue we conducted a gene enrichment analysis of all host genes. Gene enrichment analyses at each age/mating status within the WT and *bam* hypomorph genotypes infected with each *W. pipientis* variant were conducted using gseGO, a function within clusterProfiler (27).

Of the top 10 GO categories shown in our analysis, similarities between WT and *bam* hypomorph gene enrichment include the enrichment of egg chorion genes in *w*Mel2a infected three-day old flies (Fig 5). Six-day old flies were highly enriched for genes involved in egg chorion production and female gamete generation in both the WT and *bam* hypomorph (Fig 5). Mated three-day old WT flies infected with *w*Mel3, *w*MelCS2a, or *w*MelCS2b have no genes enriched under our filters (p < 0.05, absolute log2fold change > 1, Fig 5). However, by refining our filter to genes with a p-value of less than 0.05, while excluding a log2fold change cutoff, we observed a significant enrichment of genes involved in metabolic processes in WT mated three-day old flies infected with each *W. pipientis* variant. (S3 Fig). Notably, six-day old WT gene enrichment for genes involved in eggshell formation and other oogenesis-related GO categories are remarkably similar between flies infected with each *W. pipientis* variant (Fig 5E).

**Fig 5.**
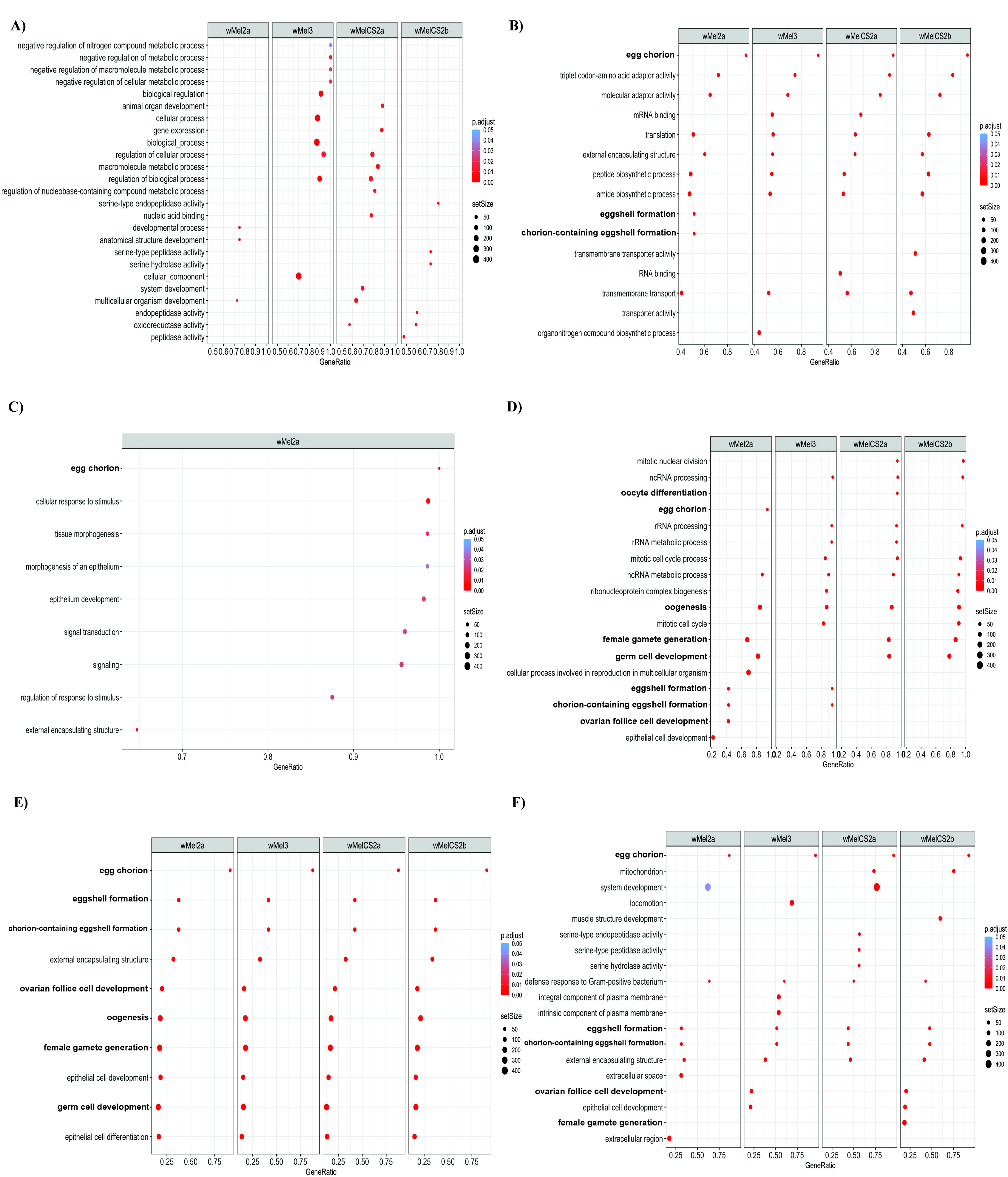
RNA-seq WT and *bam* hypomorph gene enrichment by age of female flies (p < 0.05, abs log2fold change > 1). The top 10-15 GO terms from a GO enrichment analysis of A) WT unmated three-day old flies *bam* hypomorph unmated three-day old flies, C) WT mated three-day old flies, D) *bam* hypomorph mated three-day old flies, E) WT mated six-day old flies, and F) *bam* hypomorph mated six-day old flies (gseGO, R). Each column represents the *W. pipientis* variant infecting *D. melanogaster*. The set size represents the number of genes within each GO category. The gene ratio is the number of genes within the GO term (set size) divided by the total number of differentially expressed genes. GO categories related to reproduction are highlighted in bold.

In contrast to WT gene enrichment, genes involved in reproduction were primarily enriched in mated three-day old *bam* hypomorph flies, prior to peak fertility rescue (Fig 1B and Fig 5). *Bam* hypomorph infection by *W. pipientis* variants was associated with enrichment of over 100 genes involved in oogenesis, germ cell development, female gamete generation and other oocyte-specific GO terms in mated three-day old flies (Fig 5D). While enrichment of genes involved in follicle cell development and female gamete generate occurred in six-day old *bam* hypomorph flies, they were only enriched in *w*Mel3 and *w*MelCS2b infected flies (Fig 5F). The most similar gene enrichment between *W. pipientis* variants infecting the *bam* hypomorph occurred in unmated three-day old flies (Fig 5B). Genes enriched in unmated three-day old *bam* hypomorph flies include egg chorion related genes, genes involved in RNA processes and other biosynthetic processes (Fig 5B).

### *W. pipientis* variant-specific gene enrichment shows suppression of genes involved in oogenesis in the *w*Mel-like infected *bam* hypomorph

Fertility differed between *bam* hypomorph flies infected with either *w*Mel-like variants or *w*MelCS-like variants, prompting us to test whether host gene expression differs among flies infected with either *W. pipientis* group. We subsetted the genes solely differentially expressed in *bam* hypomorph flies infected with *w*Mel-like (*w*Mel2a and *w*Mel3) variants or *w*MelCS-like variants (*w*MelCS2a and *w*MelCS2b). Gene enrichment analysis of mated three-day old *bam* hypomorph flies infected with *w*Mel-like *W. pipientis* variants showed downregulation of genes involved in egg formation and oogenesis-related GO categories (Fig 6B). Gene enrichment of oogenesis-related genes does not occur in WT flies infected with *w*Mel-like or *w*MelCS-like variants at any age (S6 Fig). Genes associated with *w*Mel-like specific GO categories related to reproduction (as depicted in Fig 6B) in mated three-day-old *bam* hypomorph flies, play roles in egg chorion formation (S4 Table). In the WT genotype, these genes are upregulated or are not differentially expressed (S4 Table). There were no signatures of enrichment of oogenesis-related genes in flies infected with *w*Mel-like or *w*MelCS-like variants in unmated three-day old or mated six-day old *bam* hypomorph flies (Fig 6A and 6C).

**Fig 6.**
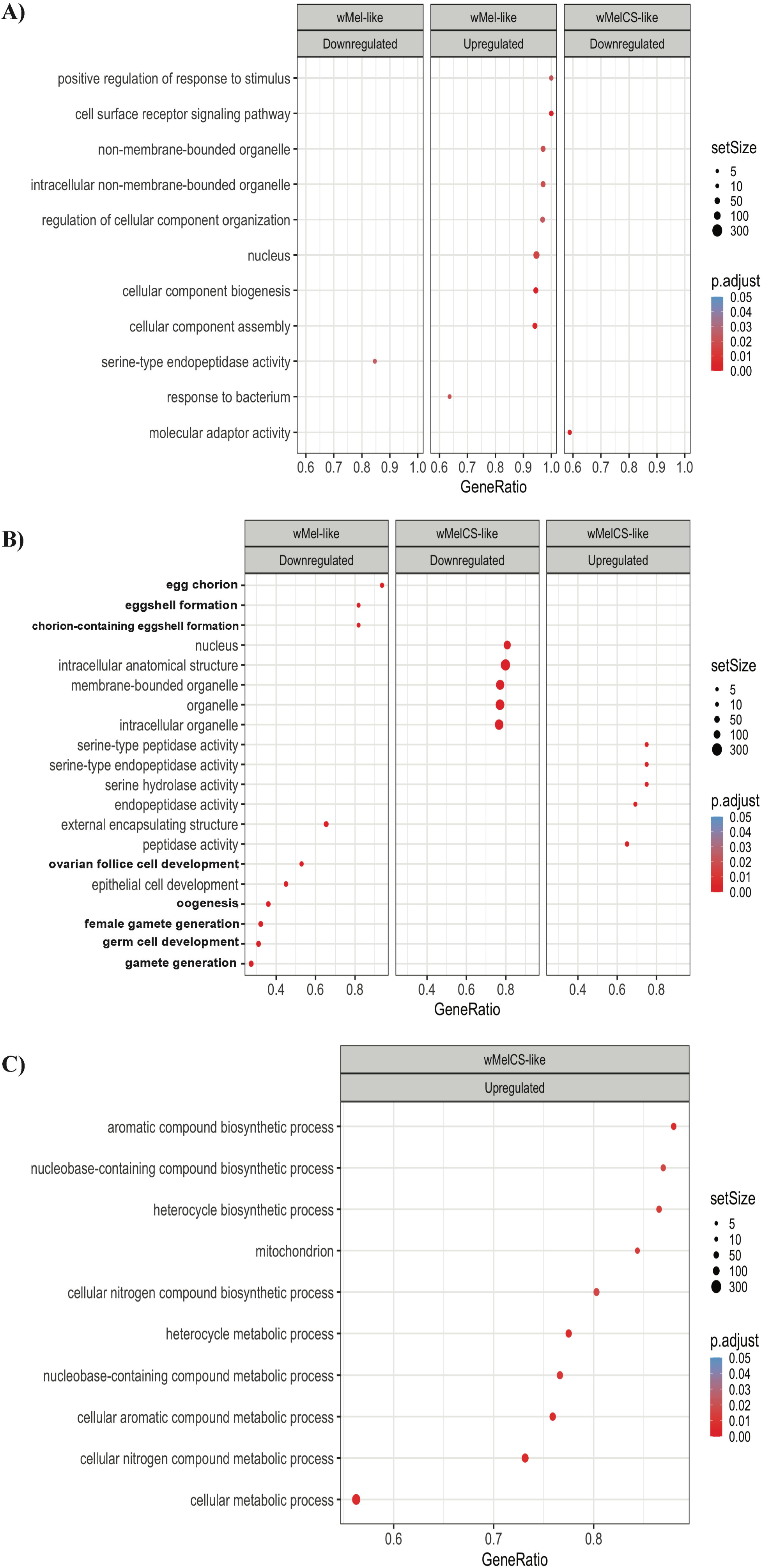
*W. pipientis* infection-specific *bam* hypomorph gene enrichment (p < 0.05, abs log2fold change > 1). A GO enrichment analysis of genes up or downregulated in each *W. pipientis* group (*w*Mel-like and *w*MelCS-like) infecting the *bam* hypomorph. A) Unmated three day-old *w*Mel-like, B) unmated three-day old *wMelCS*-like, C) mated three-day old *w*Mel-like, D) mated three-day old *w*MelCS-like, E) mated six-day old *w*Mel-like, F) and mated six-day old *w*Mel-like infected *D. melanogaster* gene enrichment. The set size represents the number of genes within each GO category. The gene ratio is the number of genes within the GO term (set size) divided by the total number of differentially expressed genes. GO categories related to reproduction are highlighted in bold.

## Discussion

Our RNA-seq analysis uncovered a subset of genes known to be involved in cystoblast differentiation that could be associated with the *bam* hypomorph fertility rescue. The task now becomes understanding the extent to which these GSC gene expression differences in *Drosophila* play a role in the observed increase in female *bam* hypomorph fertility caused by *W. pipientis* infection. Because *W. pipientis* is used as a biological control for vector-borne diseases by manipulating reproduction in the host, it is of particular interest and importance to understand how *W. pipientis* mechanistically interacts with host GSC genes[18, 19].

Our RNA-seq analysis is the first to compare gene expression profiles in *D. melanogaster* infected with four *W. pipientis* variants representing two major types (*w*Mel-like and *w*MelCS-like). Thousands of genes were differentially expressed in three-day old unmated and three- and six-day old, mated WT and *bam* hypomorph flies infected with four *W. pipientis* variants, showing the robustness of our dataset (S1 File). While it is difficult to compare our dataset with others, our unmated three-day old WT data is most comparable to the RNA-seq study of He et al. [15] that used 80 pairs of ovaries per replicate dissected from four-day old virgin females infected and uninfected with the *w*Mel strain of *W. pipientis* [20]. The GO categories that are enriched in our unmated three-day old *w*Mel2a and *w*Mel3 infected WT flies and in their dataset include: macromolecule metabolic process, oxidoreductase activity, and developmental process [20]. The He et al (15) study did not use Bonferroni correction for multiple hypothesis testing or a log2fold change cutoff for their gene enrichment analyses as we did for our analyses shown in figure 5. Therefore, we removed multiple hypothesis testing and the log2fold change filter from our gene enrichment analysis for comparison (S4 Fig). From all the enriched GO categories in this reanalysis (only the top 10 are shown in figure 5), there are 25 GO categories that are enriched in both datasets [20] supporting the integrity of our data compared to this other dataset (S1 file, S4 Fig).

We hypothesized that the differentially expressed genes that are specific to the *bam* hypomorph rather than the WT are more likely to be involved in the *bam* hypomorph fertility rescue by *W. pipientis.* Therefore, we were most intrigued by *bam’s* physical interactors and other GSC genes that were differentially expressed in mated three-day old *bam* hypomorph flies that had the largest magnitude of expression changes at a given age/mating status (Fig 4). For example, *bgcn* was highly expressed in the *bam* hypomorph (log2fold change > 5) and is of particular interest to us because its protein directly binds to *bam* in WT female flies to promote GSC daughter cell differentiation into cystoblast (Fig 4B, [21]).

In the GSC daughter cell differentiation GO category in *bam* hypomorph mated three-day old flies, three genes (*ubi-p63E*, *otu*, and *cycA*) are known to be involved in ubiquitination and deubiquitination [22]. In the WT genotype, *ubi-p63E*, *otu*, and *cycA* are not highly differentially expressed (Fig 4B). Otu is a protein that deubiquitinates CycA, and this action promotes the differentiation of GSC daughter cells into cystoblasts. [22]. Ubi-p63E is specifically associated with a distinct phenotype observed exclusively in males. Ubi-p63E protein supplies free ubiquitin and plays an essential role in spermatid differentiation [23–26]. As free ubiquitin is needed for CycA turnover, it is possible the Ubi-p63E plays a similar role in females as it does in males considering that it is highly differentially expressed in our analysis (Fig 4B). *Apc2* which is only differentially expressed in *w*Mel-like infected *bam* hypomorph flies, is a ubiquitin ligase known to interact with CycA to promote proper egg chamber formation (S4 Table, [24–26]. While *cycA* was computationally predicted to be influenced by tissue size in the three-day old *bam* hypomorph flies, it is possible that expression of *cycA* could be influenced by both tissue size and *W. pipientis* infection. Interestingly, genes associated with deubiquitination are known to trigger cytoplasmic incompatibility in male insects infected with *W. pipientis* [27, 28], underscoring the wider significance of our research findings. The differential expression of these ubiquitin-related genes in the *bam* hypomorph leads us to believe that ubiquitination and deubiquitination could be important process for the *bam* hypomorph fertility rescue by *W. pipientis* to occur.

Genes involved in histone lysine methylation are also enriched in mated three-day old *bam* hypomorph flies (Figure 4B). Histone H3K9 trimethylation specifically is known to be involved in GSC differentiation and maintenance in *D. melanogaster* [14, 15, 17]. Three differentially expressed genes specific to the *bam* hypomorph in the histone lysine methylation category, *wde, egg, and su(var)3-9*, have known functions in GSC differentiation or maintenance [15–17]. Moreover, the fact that host methylation, such as the activity of the DNA methyltransferase Dnmt2, has been found to influence pathogen blocking in *D. melanogaster* males, further underscores the overarching significance of methylation in various processes in *D. melanogaster*, including fertility rescue in *bam* hypomorphs [29–31].

The COP9 complex is disrupted by Bam to inhibit GSC self-renewal [32]. Our analysis shows that it is unlikely that genes in the COP9 complex are associated with the *bam* hypomorph fertility rescue at the transcriptional level because they are not highly differentially expressed (p < 0.05, abs log2fold change > 1, S2 Table and S2 Fig). Additionally, genes involved in the CCR4-NOT complex are not differentially expressed in the *bam* hypomorph (S2 Fig). For GSC self-renewal inhibition to occur, a complex consisting of Bam, Bgcn, Mei-P26, Brat, a deadenylate and Ago1 or Sxl form to destabilize *nos* mRNA [21, 33, 34]. It is known that Bam binds to CAF40, a subunit of the CCR4-NOT deadenylase [35]. It is hypothesized that the CCR4-NOT deadenylase functions to destabilize *nos* mRNA and inhibit GSC self-renewal [35, 36]. The absence of differentially expressed genes involved in the COP9 complex and the CCR4-NOT complex in the *bam* hypomorph leads us to believe that the *bam* hypomorph fertility rescue could be independent of genes involved in GSC-self renewal. Genes in the COP9 signalosome and CCR4-NOT complex are not differentially expressed in the WT genotype, suggesting these genes are not impacted by *W. pipientis* infection (Fig 4B and S2 Table). Our hypothesis is supported by our *bam* hypomorph-specific data showing differential expression of genes involved primarily in GSC daughter cell differentiation.

In WT flies, enrichment of oogenesis-related genes in six-day old flies occurs during the greatest increase in fertility (Figs 1 and 5E). Oogenesis-related gene enrichment is surprisingly similar in flies infected with different *W. pipientis* variants in the WT genotype, despite the difference in fertility between *w*Mel-like and *w*MelCS-like infected flies (Figs 1 and 5E). The *w*Mel-like and *w*MelCS-like variants of *W. pipientis* exhibit distinct phenotypes in *Drosophila melanogaster*, encompassing variations in viral protection, fecundity, and cytoplasmic incompatibility. While we hypothesized that the difference in fertility between WT *w*Mel-like versus *w*MelCS-like infected flies could be due to *W. pipientis* variant-specific infected gene enrichment, our analysis did not find any oogenesis-related gene enrichment in either *W. pipientis* group (S5 Fig). The host could have developed compensatory mechanisms to counteract the differences in gene expression caused by *W. pipientis* variants or oogenesis-related genes could be selected for among flies infected with any *W. pipientis* variant in order to maintain proper development.

Gene enrichment analysis of oogenesis-related genes in three-day-old *bam* hypomorph flies, prior to peak fertility rescue, revealed variation between flies infected with different *W. pipientis* variants (Fig 5B). This variation in gene enrichment could potentially explain the observed differences in fertility rescue efficiency between flies infected with different *W. pipientis* variants (Fig 1B). Given that germline stem cells require several days to undergo differentiation into mature eggs, the observed differential expression of oogenesis-related genes prior to peak fertility rescue in the *bam* hypomorph is not unexpected [37, 38]. Our enrichment analysis of the *W. pipientis* variant specific genes showed *w*Mel-like infected *bam* hypomorph flies downregulate genes involved in egg chorion production and five genes involved in either oogenesis or spermatogenesis (S4 Table). *Apc2* downregulation promotes germarium-derived egg chamber formation [25]. It is possible that *w*Mel-like infected flies downregulate egg chorion genes as an immune response to *W. pipientis*, but downregulation of genes such as *apc2* could potentially play a role in fertility rescue as well.

Our research sheds light on the complex interaction between host, bacterium, and reproductive system. We have identified a subset of genes involved in cystoblast differentiation and maintenance, which could contribute to the observed increase in female *bam* hypomorph fertility caused by *W. pipientis* infection. Moreover, the differential expression of ubiquitin-related genes and genes involved in histone lysine methylation suggests potential mechanistic pathways underlying the fertility rescue. Understanding the intricate relationship between *W. pipientis* variants, host GSC genes, and oogenesis-related processes will contribute to the broader understanding of host-pathogen interactions and reproductive biology.

## Methods and Methods

### Fly strains and husbandry

The four *W. pipientis* variants and uninfected control used in this study were generously provided by Luis Teixeira [7]. The generation of the *bam*^L255^ hypomorph and our control *bam w*^1118^ fly lines, which we consider as our wild-type (WT) line, containing each of the four *W. pipientis* variants, was described in the study conducted by Bubnell et al. in 2021[4]. Of note, the *bam* hypomorph mutation we use in our current study was recently remade using the same single amino acid change as the original *bam*^BW^ hypomorph but in a *w*^1118^ isogenic background [4, 39]. The female *bam* hypomorph (*w*^1118^;*bam*^L255F^/*bam*^[3xP3dsRed]^) that was used in this study was generated by crossing the *bam*^L255^/TM6 female to a *bam* null (*w*^1118^;*bam*^[3xP3dsRed]^ /TM6) male [4]. Additionally, an uninfected *bam* hypomorph control was used along with WT *bam w*^1118^ fly lines containing each *W. pipientis* variant.

All *D. melanogaster* lines were maintained on yeast glucose food and placed in an incubator at 25°C with a 12-hour light-dark cycle.

### W. pipientis variants

All experiments were conducted with one of four *W. pipientis* variants (*w*Mel2a, *w*Mel3, *w*MelCS2a and *w*MelCS2b) infected with *D. melanogaster* and an uninfected control in the WT and *bam* hypomorph genotypes. *W*Mel2a and *w*Mel3 are referred to as *w*Mel-like variants while *w*MelCS2a and *w*MelCS2b are referred to as the *w*MelCS-like variants. PCR was used to confirm that the correct *W. pipientis* variant infected *D. melanogaster* using *W. pipientis*-specific primers listed in Riegler et al. 2005 [8].

### Developmental time course and time interval fertility assay

Aged virgin WT or *bam* hypomorph females were mated with two to three-day old Canton-S males for 24 hours. We used unmated day three-day old, mated three-day old, mated six-day old, and mated nine-day old female WT and *bam* hypomorph females. 10 WT or *bam* hypomorph female flies and 10 Canton-S males were mated in a single vial with nine additional replicates per age and mating status. For example, 10 two-day old *bam* hypomorph females were mated with 10 two- to three-day old Canton-S males at 11 AM and collected at 11 AM the next morning.

We collected the female parents of different ages/mating statuses to measure *W. pipientis* titer and conduct RNA-seq. Subsequently, a time interval fertility assay was conducted by measuring the number of progeny from each vial in which the parents mated. The progeny per vial were counted every two days for eight days and the sum of progeny was used for further analysis.

A Poisson response distribution was conducted in R (v. 4.1.0) at each timepoint to determine statistically significant differences in fertility between *D. melanogaster* infected with different *W. pipientis* groups (*w*Mel-like and *w*MelCS-like) and compared to the uninfected control. The equation used was as follows: Fertility (response variable) ∼ *W. pipientis* Group * Day.

### Ovary dissections and DNA/RNA extractions

Ovaries of the parent female flies from the time interval assay were dissected in cold PBS using two forceps. Zymo’s Quick-RNA 96 kit was used to extract DNA and RNA from dissected female parent ovaries.

### *W. pipientis* titer quantification

QPCR was used on the ovarian parent fly DNA from the time interval fertility assays to quantify *W. pipientis* titer. Absolute quantification was used to quantify *W. pipientis* titer. Absolute quantification compares DNA of an unknown quantity to a standard curve made from DNA of a known quantity. This method was shown to be more efficient in measuring *W. pipientis* titer than relative quantification [40]. We created a gBlock containing *W. pipientis* DNA fragments (S6 Fig) and used the *W. pipientis*-specific gene *DNA recombination-mediator protein A*, *dprA*, for our standard curves. QPCR was run on a QuantStudio 7 Pro provided by Cornell’s Biotechnology Resource Center. A linear regression model was conducted in R (v. 4.1.0) at each timepoint to determine statistically significant differences in *W. pipientis* titer between *D. melanogaster* infected with different *W. pipientis* groups (*w*Mel-like and *w*MelCS-like) and between the uninfected control. The equation used was as follows: *W. pipientis* titer (response variable) ∼ *W. pipientis* Group * Day.

To assess correlations between *D. melanogaster* fertility and *W. pipientis* titer, we used a linear regression model. The equation used was as follows: *D. melanogaster* Fertility ∼ *W. pipientis* Titer.

### Choosing timepoints and biological replicates for RNA-seq

The time interval fertility assays showed that peak fertility rescue occurred in six-day old flies. Therefore, six-day old female flies were used for RNA-seq. Additionally, we used unmated three-day old female flies as a mating control and mated three-day old female flies as an age control for RNA-seq. Four biological replicates were used for each age/mating status.

### Library preparation and 3’ RNA sequencing

Ovaries from *bam* hypomorph and WT flies infected with each *W. pipientis* variant, and the uninfected controls were dissected, DNA/RNA extractions were conducted, and the RNA was subsequently used for 3’RNA seq library construction and Illumina NextSeq500 (conducted by the Cornell Genomics Facility).

### Differential expression analysis

RNA-seq reads were mapped to the reference *D. melanogaster* genome (Flybase) using STAR [41]. FastQC of the FastQ files showed that quality scores were high (>30 phred score), and adapter content was low (<2% of sequence). Therefore, no reads were trimmed. FastQC of the aligned sequences showed that sample alignment percentage on average was 87%. However, one WT three-day old biological sample infected with *w*Mel3 aligned below 70% and was removed. A read count matrix was made and used for differential expression analysis using DESeq2 [42].

WT ovaries are significantly bigger than *bam* hypomorph ovaries, even when rescue by *W. pipientis* occurs. WT ovaries also consistently contain differentiated cystoblasts while *bam* hypomorph ovaries only contain some differentiated cystoblasts when rescue of the tumorous phenotype occurs, but some GSC-like undifferentiated cells remain. Considering that there are significant differences between the tissue size and type of *bam* hypomorph and WT ovaries, we could not compare expression between these genotypes directly. Therefore, differential expression analyses compared *W. pipientis* infected *D. melanogaster* expression relative to uninfected *D. melanogaster* expression within genotypes. The differential expression within genotypes was then compared between genotypes. Volcano plots showing the total amount of genes up and downregulated at each age/mating status in the *bam* hypomorph and WT genotypes infected with each *W. pipientis* variants were made using R (S1 File).

### Tissue size, mating, and age analyses

Genes influenced by tissue size were identified computationally using glm.nb. Specifically, a null formula containing expression as the dependent variable and infection status as an independent variable was compared to a test formula containing expression as the dependent variable, and infection status and tissue size as independent variables. A ranking for *W. pipientis* rescue in the *bam* hypomorph determined from the time interval fertility assay was used as a proxy for tissue size because tissue size increases as the rescue of the fertility phenotype increases.

Glm.nb, a function in the MASS R package, was used to identify genes impacted by age and mating. The null formula included expression as the dependent variable and the *W. pipientis* variant as the independent variable. The test formula included expression as the dependent variable along with *W. pipientis* variant and age/mating as the independent variables. Genes impacted by age, mating, or tissue size were then identified and noted in the supplement (S4 Table). However, we did not remove the genes from the age, mating, and tissue size analysis from our dataset as it is possible that these genes could both be impacted by one of the factors and still impact *bam* hypomorph fertility rescue.

### GO analyses

GO analyses were conducted using gseGO within the clusterProfiler package in R (v.1.4. 1717). Differentially expressed genes with a p-value above 0.05 were removed prior to the analysis. The genes were then ranked by log2fold change prior to using gseGO. Our gseGO analysis included all gene ontologies (biological process, cellular component, and molecular function).

## Acknowledgements

We thank Andy Moeller and Andy Clark for insightful discussions and review of the manuscript. We also thank Mariana Wolfner, Jackie Bubnell, Miwa Wenzel, and Luke Arnce for thoughtful feedback throughout this project. We thank Cornell Institute of Biotechnology Imaging Center for the Zeiss i880 LSM880 microscope and the Cornell Genomics Facility for conducting 3’ RNA-seq.

---

**S1 Table. WT and *bam* hypomorph fertility assay raw data.**

The number of progeny are listed for the uninfected WT and *bam* hypomorph flies and flies infected with *w*Mel2a, *w*Mel3, *w*MelCS2a, and *w*MelCS2b. The *W. pipientis* variants were grouped into the *w*Mel-like variants (*w*Mel2a and *w*Mel3) and the *w*MelCS-like variants (*w*MelCS2a and *w*MelCS2b) in the “*Wolbachia* type” column. The parents of the progeny were either three-, six- or nine-days old.

**S1 Fig. *Bam* WT fertility does not correlate with *W. pipientis* titer.**

WT fertility and titer of all *W. pipientis* variants infecting *D. melanogaster*. Fertility and titer of mated aged flies: (A) three-, (B) six- and (C) nine-day old females. The p-value indicates whether fertility and titer are statistically correlated for each *W. pipientis* type (linear regression model).

**S2 Figure. Differential expression of *bam’s* interactors shows genes involved in ubiquitination/deubiquitination.**

Gene networks of all *bam’s* genetic and physical interactors in the A) WT unmated three-day old flies B) *bam* hypomorph unmated three-day old flies, C) WT mated three-day old flies, D) *bam* hypomorph mated three-day old flies, E) WT mated six-day old flies, and F) *bam* hypomorph mated six-day old flies (p < 0.05, abs log2fold change > 1, STRING, Cytoscape). Grey circles surrounding the genes indicate that the gene was not differentially expressed in the given *W. pipientis* infected genotype compared to the uninfected genotype.

**S2 Table. Differentially expressed *bam* genetic/physical interactors and GSC genes in the WT and *bam* hypomorph genotypes.**

B*am’s* differentially expressed genetic and physical interactors and GSC genes in WT and *bam* hypomorph unmated three-, mated three-, and mated six-day old flies (p < .05, absolute log2fold change > 1). Our differential expression analysis compared *W. pipientis* variant infected flies to uninfected flies in the WT and *bam* hypomorph genotypes. NA indicates that a given gene was not differentially expressed in each genotype infected with a specific *W. pipientis* variant.

**S3 Figure. RNA-seq WT and bam hypomorph gene enrichment by age of female flies.**

The top 10-15 GO terms from a GO enrichment analysis of A) WT unmated three-day old flies B) *bam* hypomorph unmated three-day old flies, C) WT mated three-day old flies, D) *bam* hypomorph mated three-day old flies, E) WT mated six-day old flies, and F) *bam* hypomorph mated six-day old flies (p < 0.05, gseGO, R). Each column represents the *W. pipientis* variant infecting *D. melanogaster*. The set size represents the number of genes within each GO category. The gene ratio is the number of genes within the GO term (set size) divided by the total number of differentially expressed genes. The GO categories are highlighted if they are involved in reproduction.

**S3 Table. Genes predicted to be differentially expressed due to tissue size, age or mating.**

We computationally predicted what genes could be influenced by tissue size, age or mating in mated three- and six-day old WT and *bam* hypomorph flies. Genes that are bolded are *bam’s* genetic or physical interactors and genes that are highlighted are GSC genes. Note that the genes listed in this table are not all differentially expressed in our analysis between *W. pipientis* infected and uninfected WT and *bam* hypomorph flies (S2 Table).

**S4 Fig. W. pipientis infection-specific WT gene enrichment (p < 0.05).**

A GO enrichment analysis of genes solely in each *W. pipientis* group (*w*Mel-like and *w*MelCS-like) infecting *D. melanogaster*. A) Unmated three-day old *w*Mel-like, B) unmated three-day old *w*MelCS-like, C) mated three-day old *w*Mel-like, D) mated three-day old *w*MelCS-like, E) mated six-day old *w*Mel-like, F) and mated six-day old *w*Mel-like infected *D. melanogaster* gene enrichment. The set size represents the number of genes within each GO category. The gene ratio is the number of genes within the GO term (set size) divided by the total number of differentially expressed genes.

**S4 Table. WMel-like infection-specific bam hypomorph gene enrichment (p < 0.05).**

The differentially expressed genes from oogenesis-related GO categories from our *W. pipientis* infection-specific gene enrichment analysis. The table includes the genes from our enrichment analysis that used genes with a p-value less than 0.05 and absolute log2fold change greater than 1 and from a separate enrichment analysis solely using a p-value cutoff below 0.05. The gene enrichment analysis was conducted in the *bam* hypomorph genotype infected with *w*Mel2a or *w*Mel3, but the log2fold change of the genes from our analysis is shown for both the *bam* hypomorph and WT genotypes for comparison. NA indicates that the given gene was not differentially expressed in each genotype infected with either *w*Mel2a or *w*Mel3.

**S5 Fig. *W. pipientis*-specific genes included in the gBlock used for absolute quantification of *W. pipientis* titer.**

The *W. pipientis-*specific genes included in our double stranded DNA fragment (gBlock) used for absolute quantification qPCR.

