## Supplementary figures and images for "Gene expression changes in *Drosophila melanogaster* females associated with the rescue of the *bag of marbles* (*bam*) hypomorph fertility defect by *Wolbachia pipientis*"

### Supplemental Figure 1

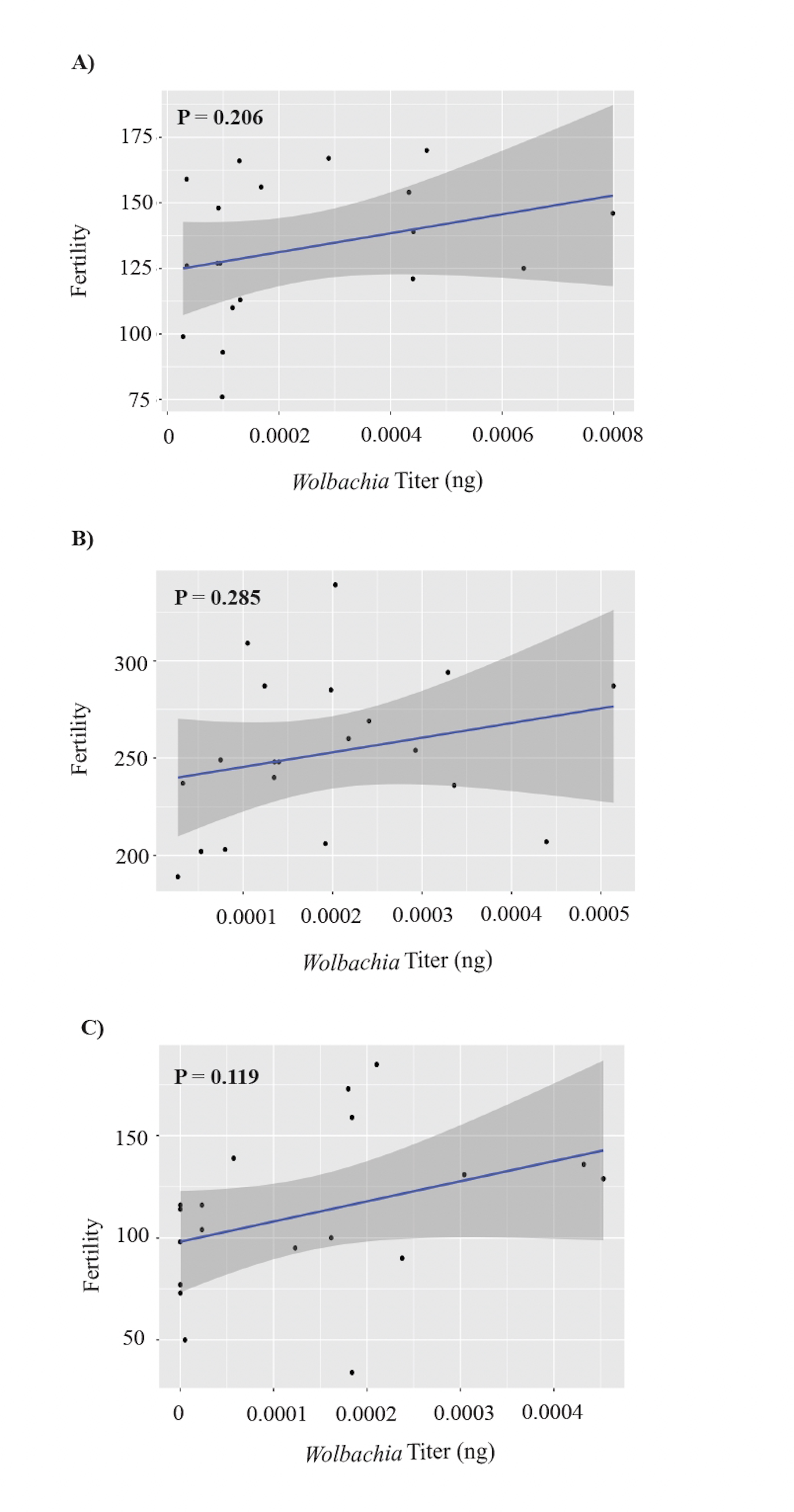

### Supplemental Figure 2

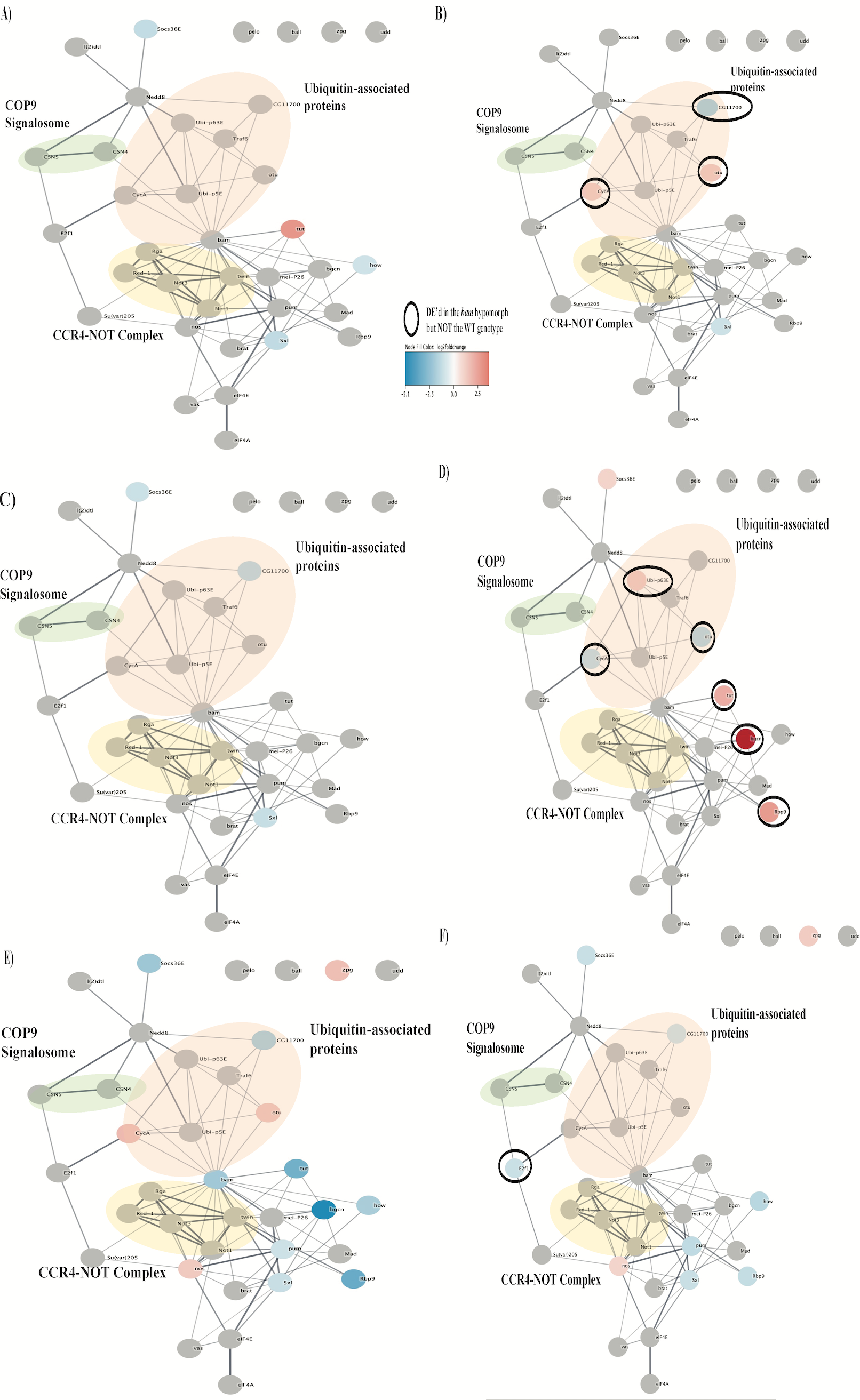

### Supplemental Figure 3

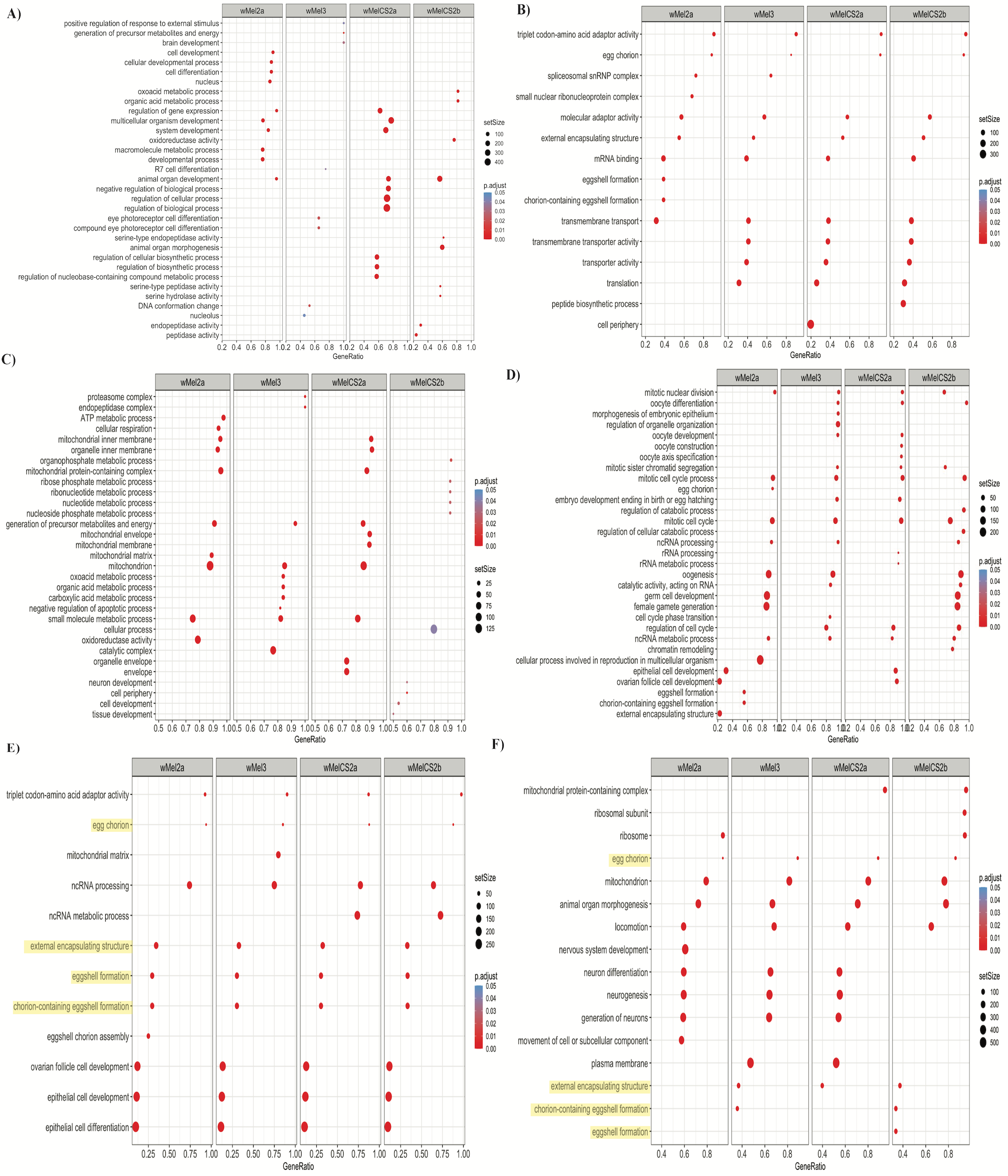

### Supplemental Figure 4

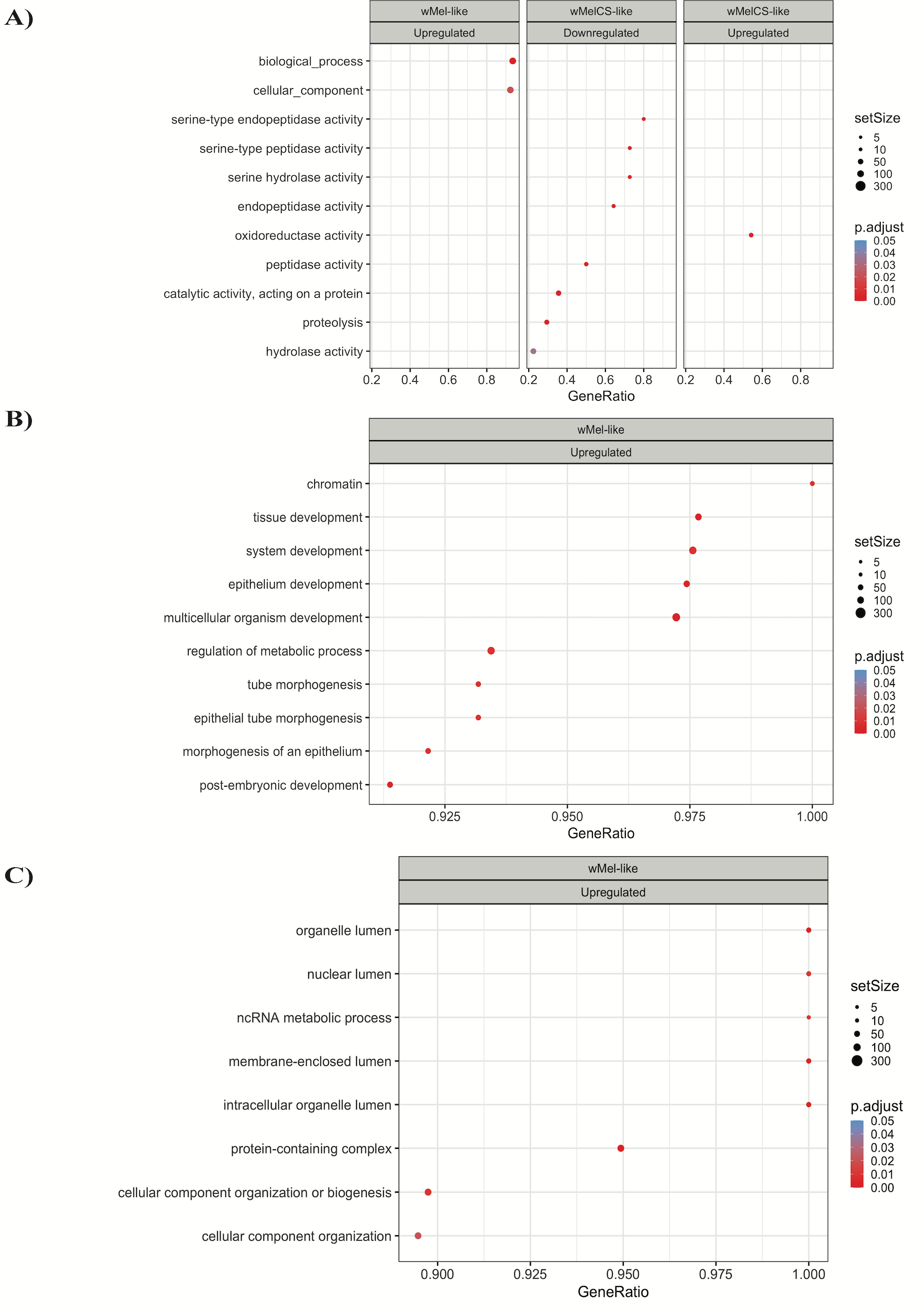

### Supplemental Figure 5

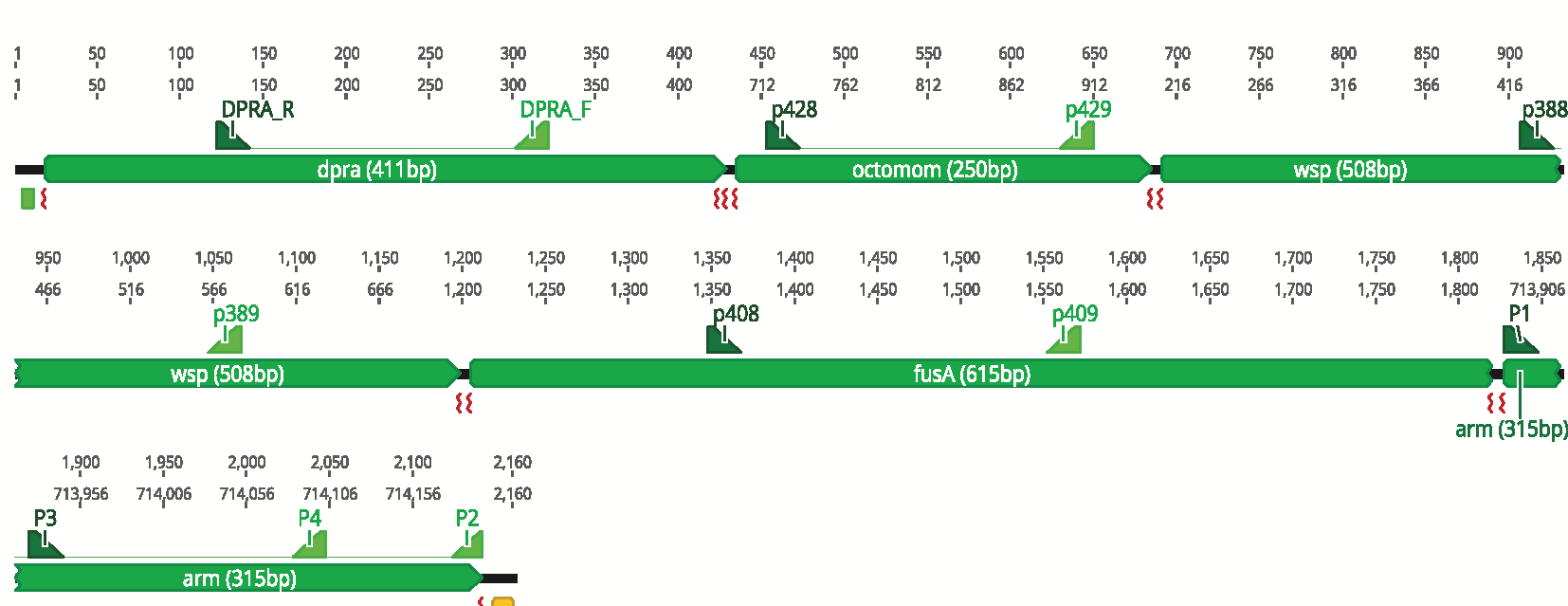
