## Supplemental File 1 for "Gene expression changes in *Drosophila melanogaster* females associated with the rescue of the *bag of marbles* (*bam*) hypomorph fertility defect by *Wolbachia pipientis*"

### Unmated WT 3 Day old

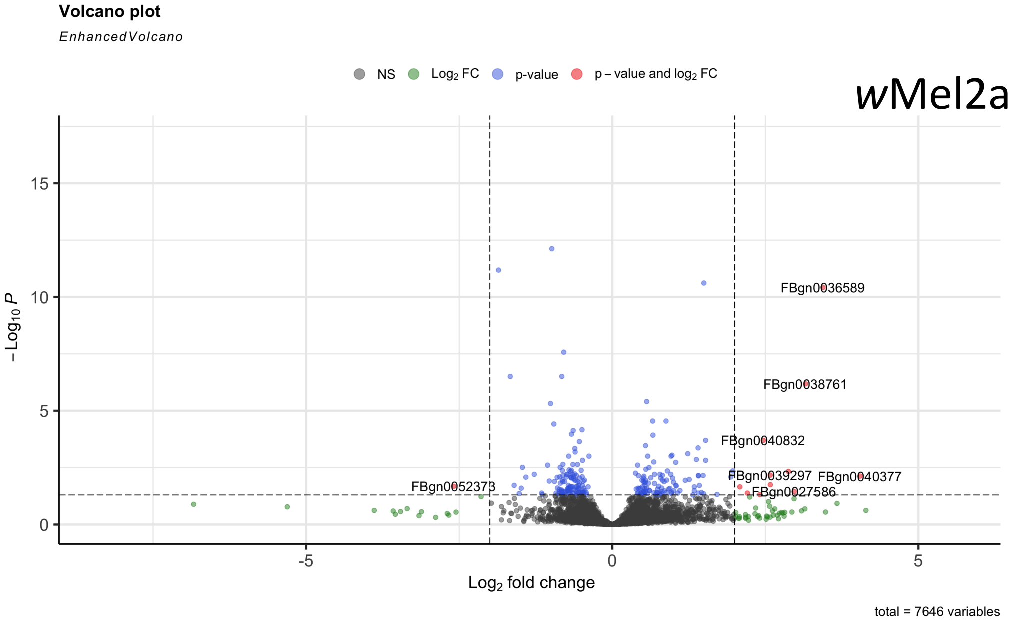


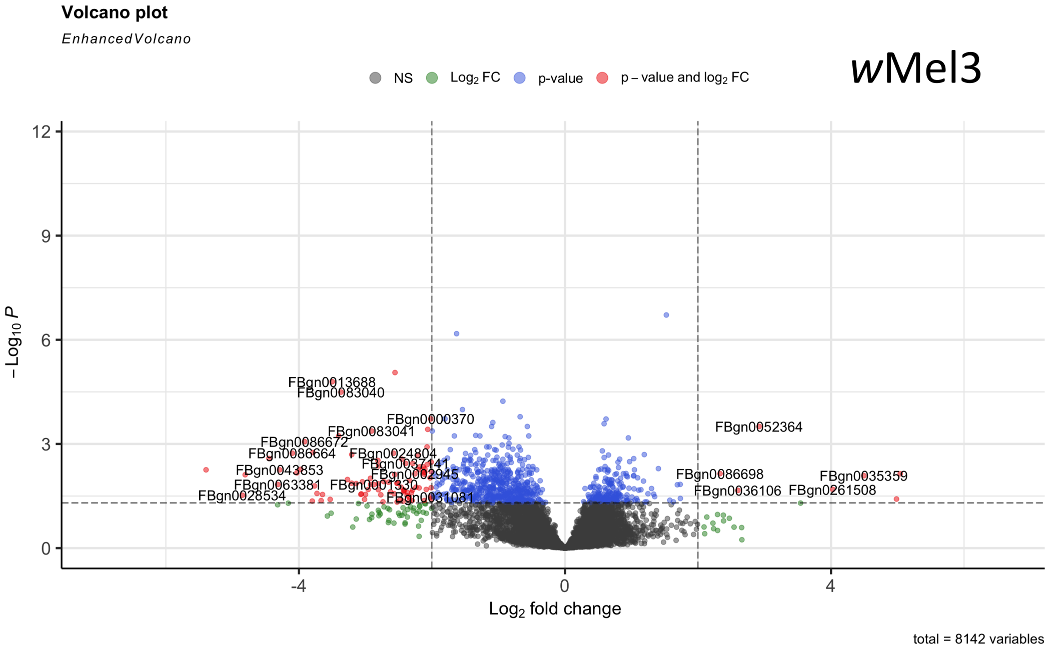


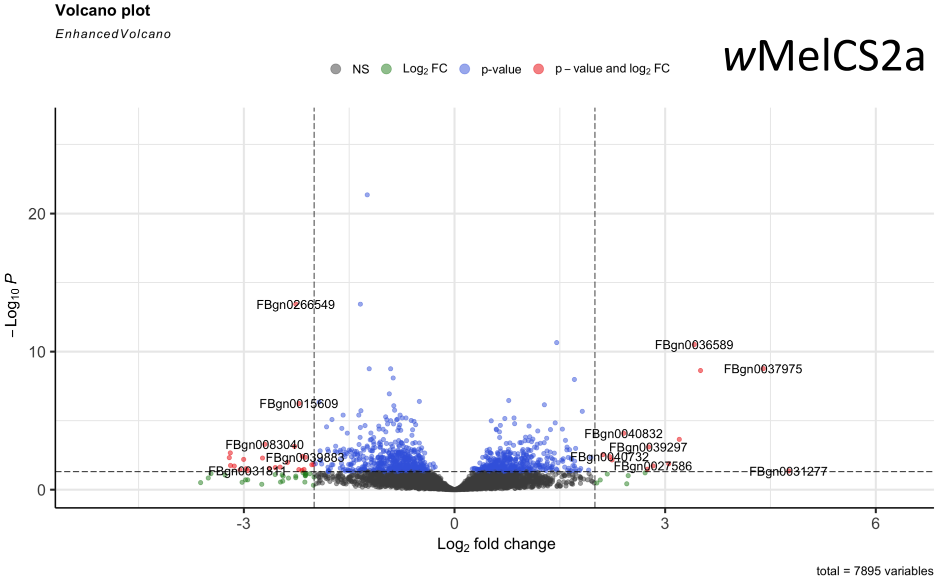


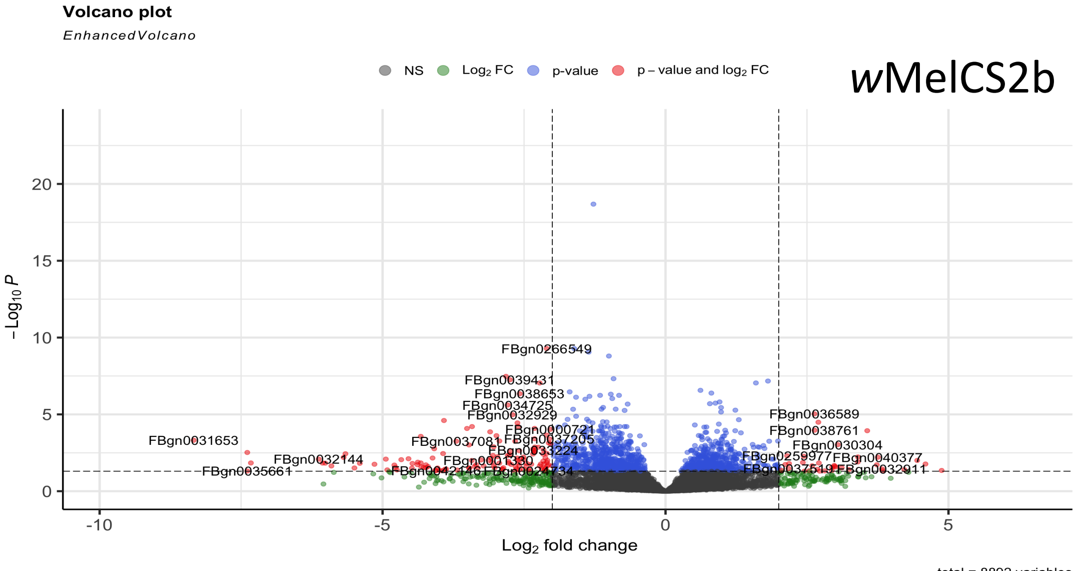

### WT Mated 3 Day old

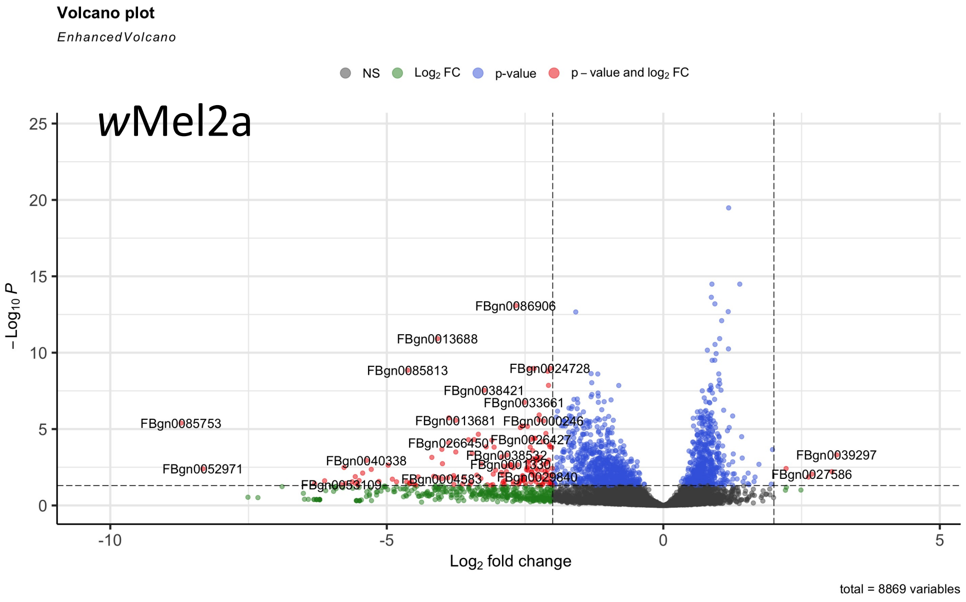


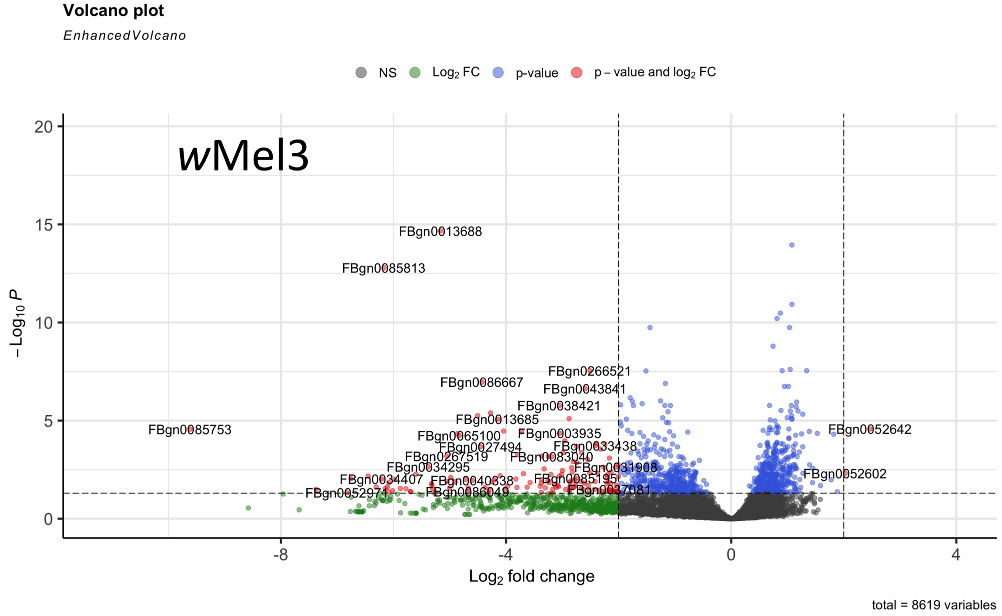


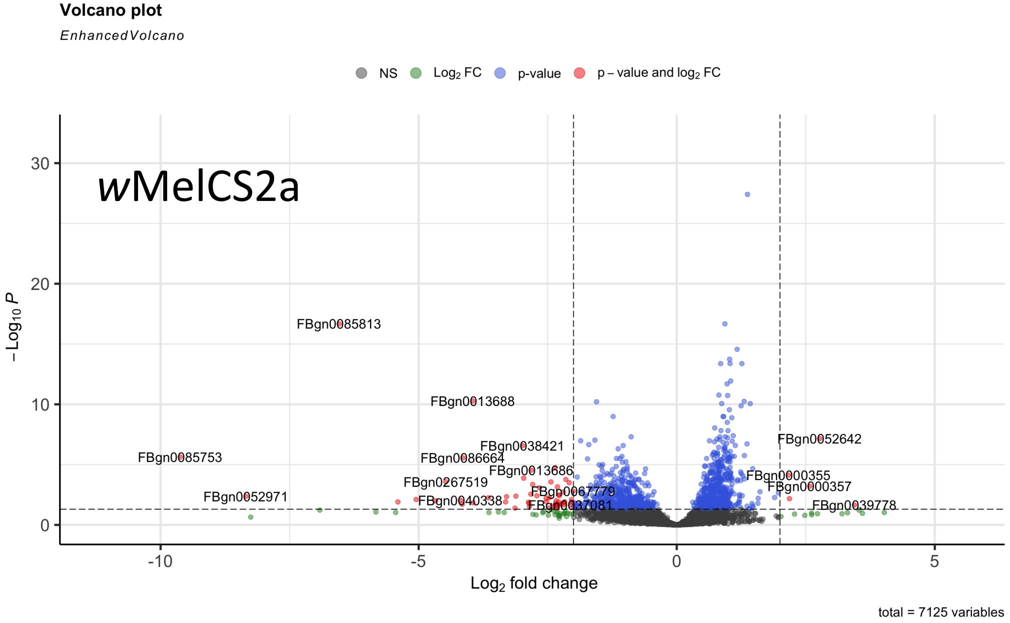


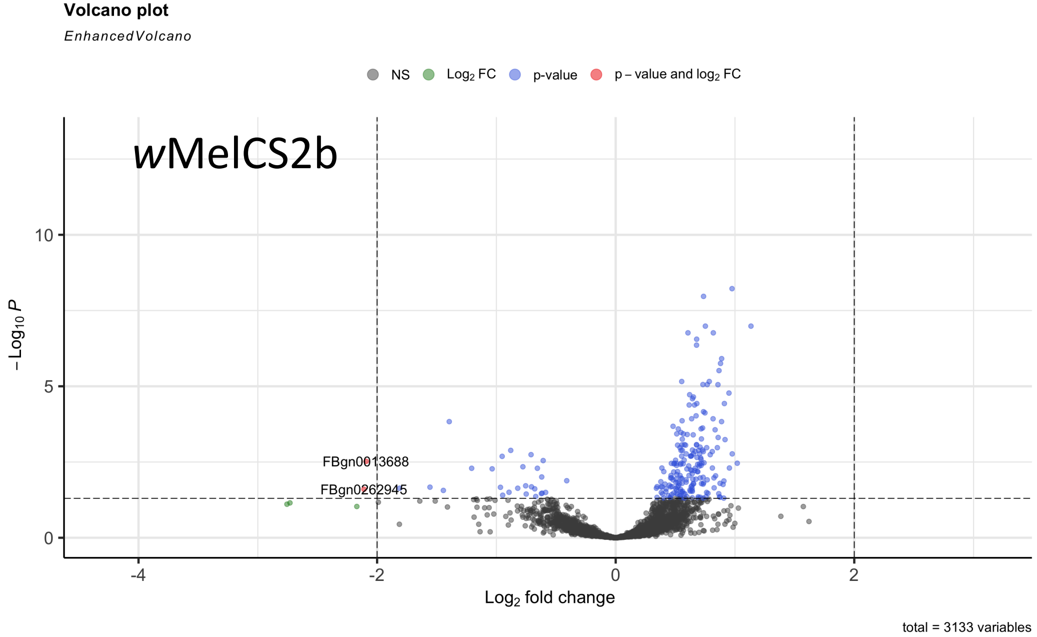

### WT Mated 6 Day old

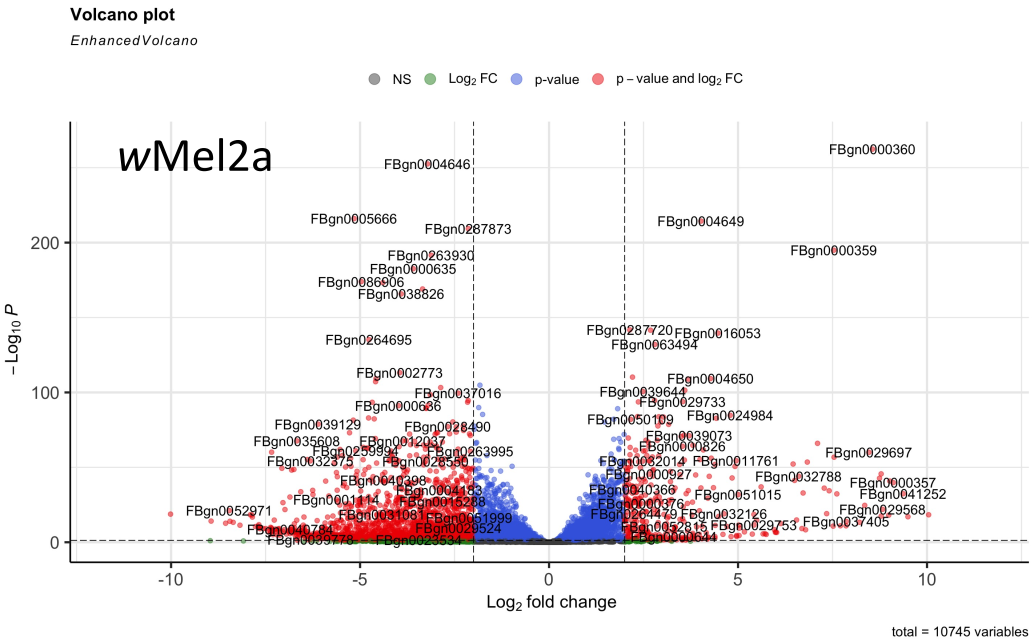


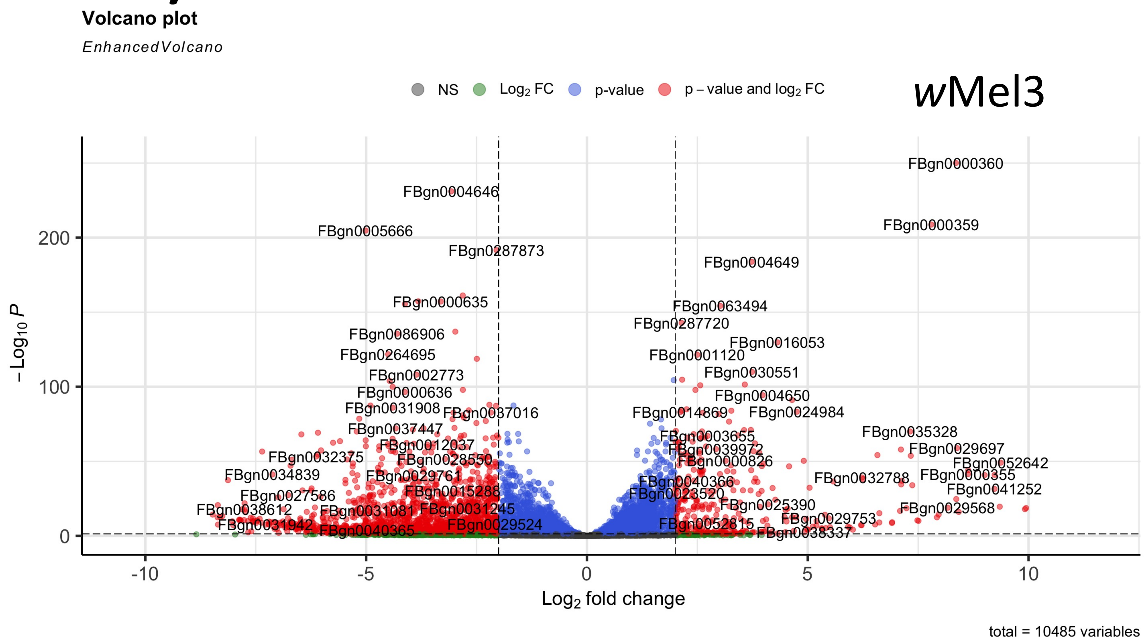


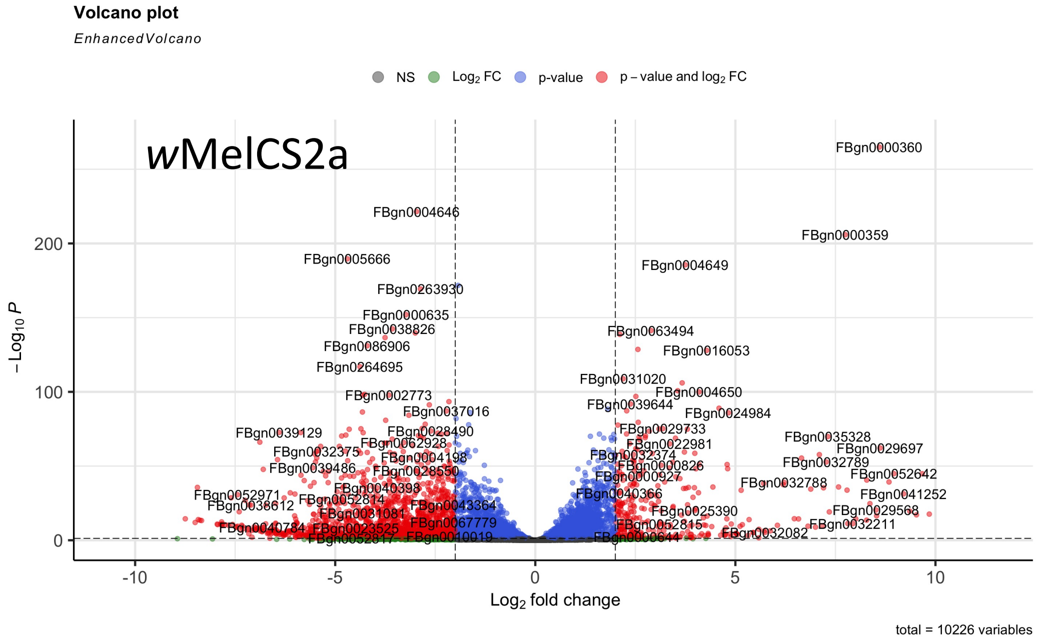


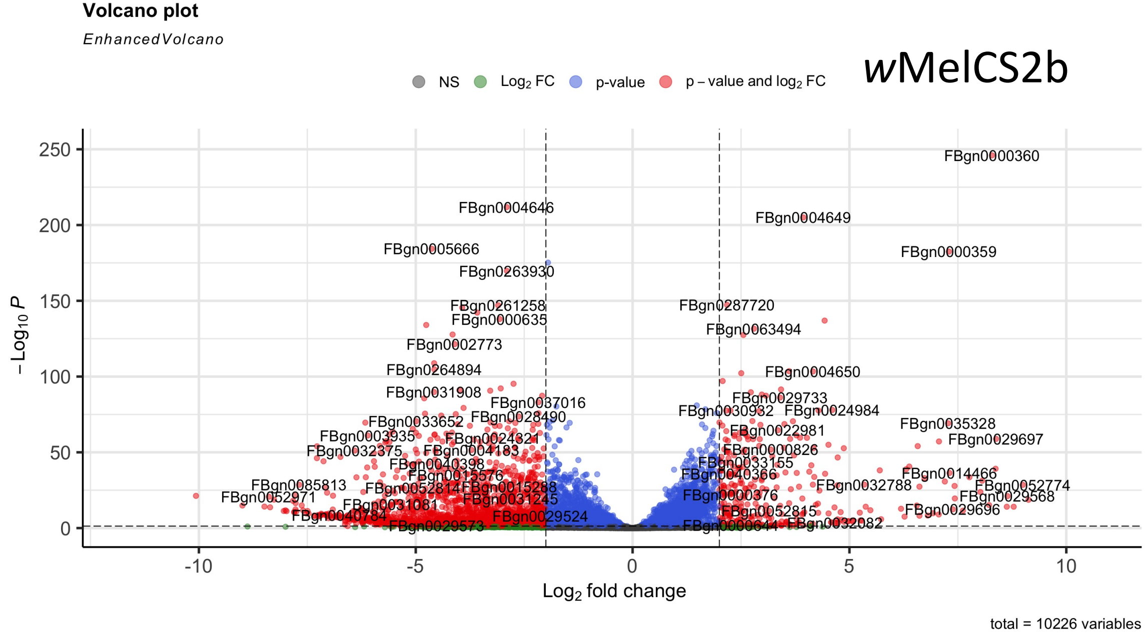

### Unmated day 3 *bam* hypomorph

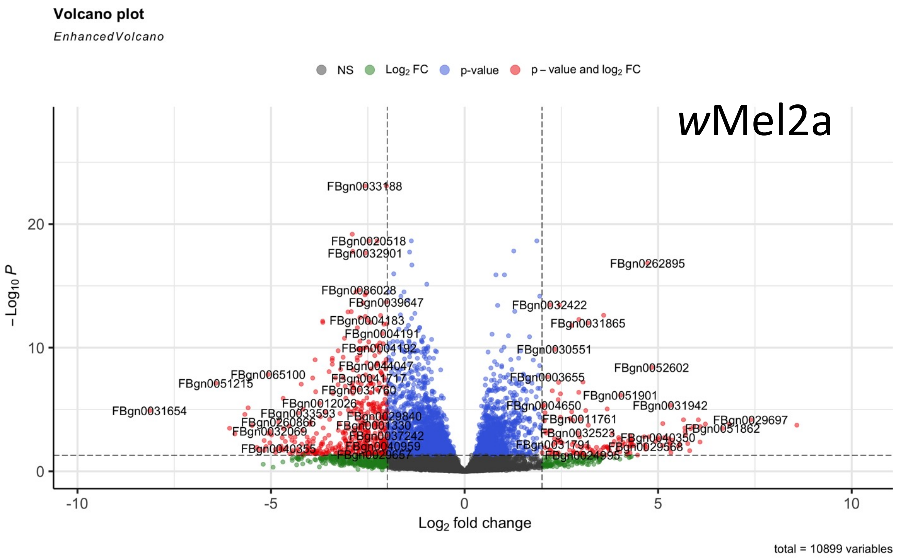


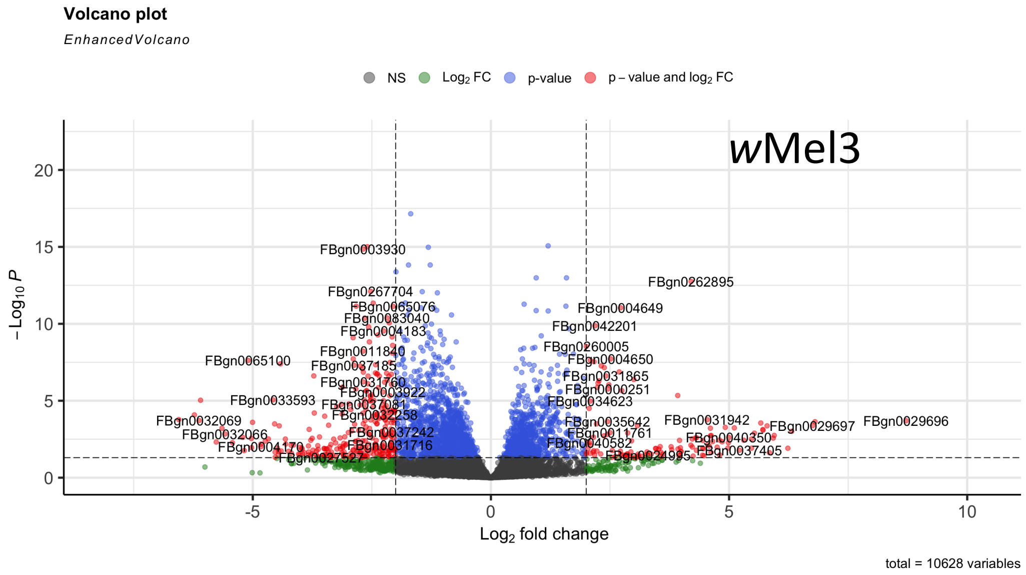


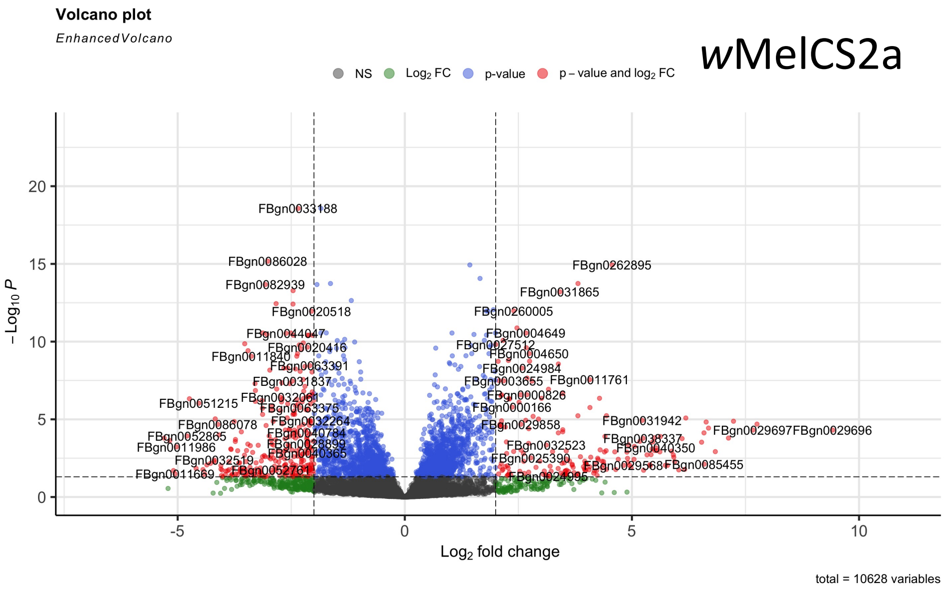


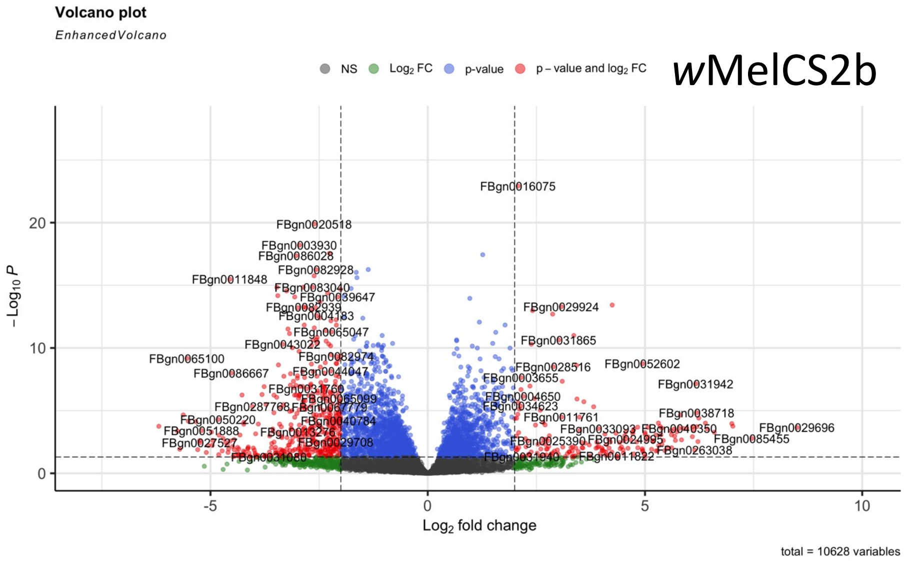

### Mated day 3 *bam* hypomorph

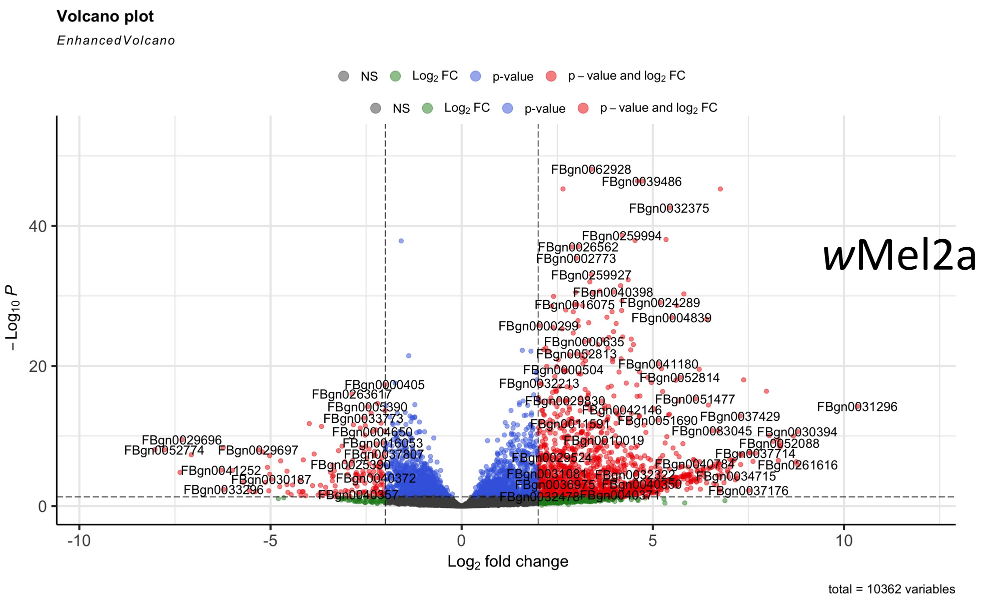


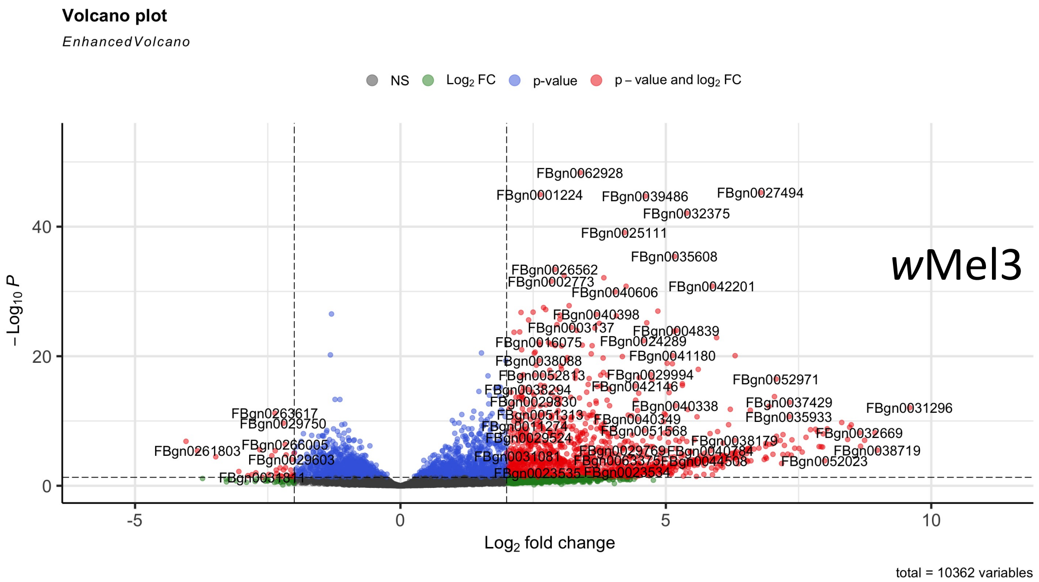


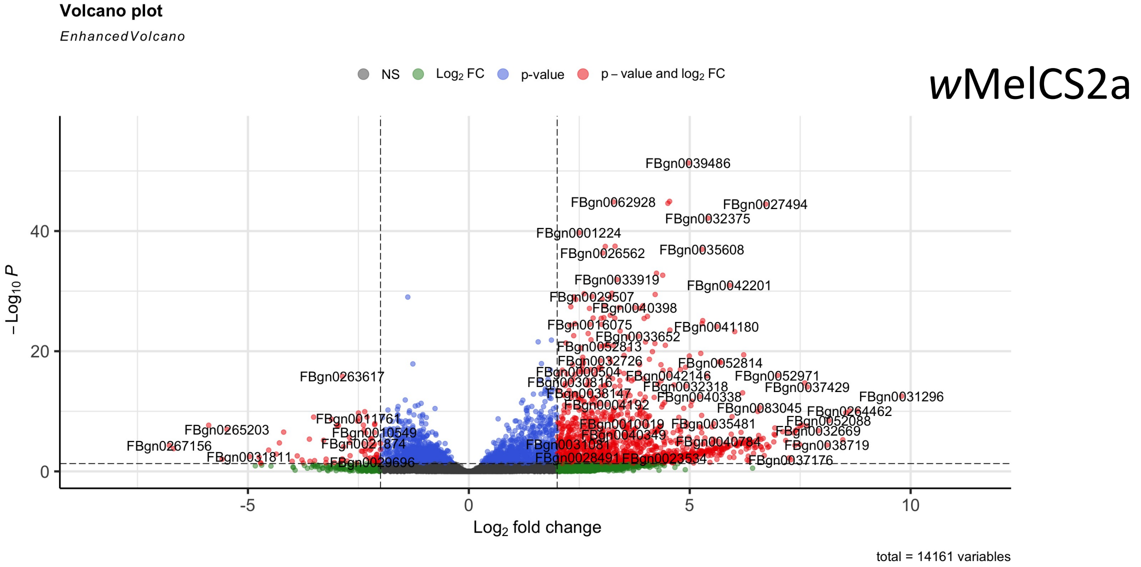


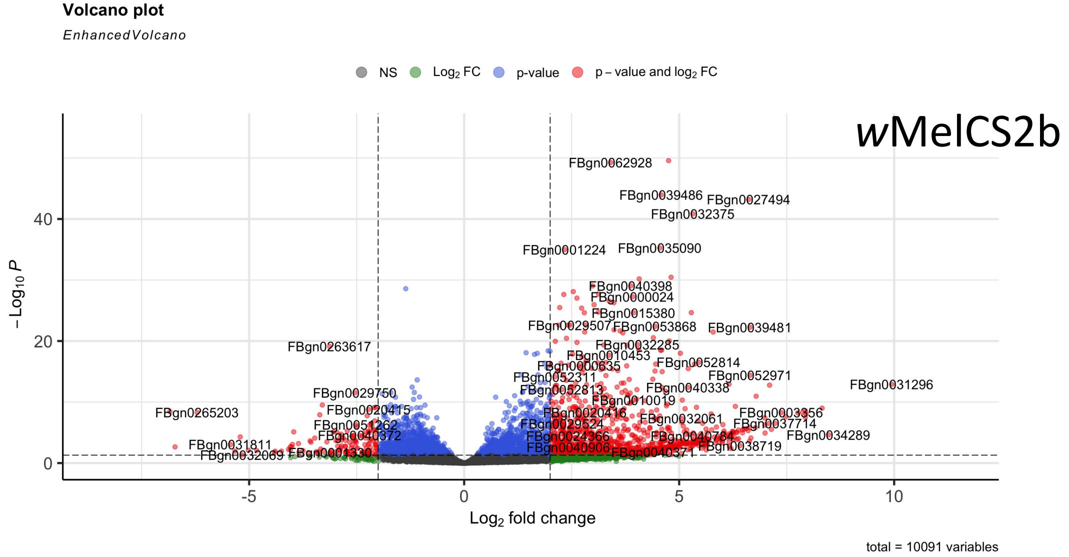

### Mated day 6 *bam* hypomorph

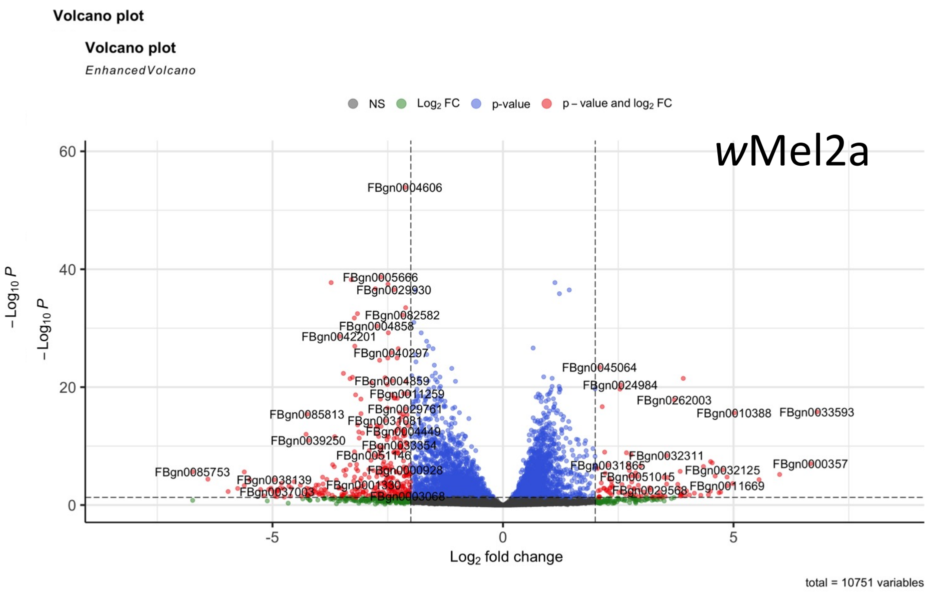


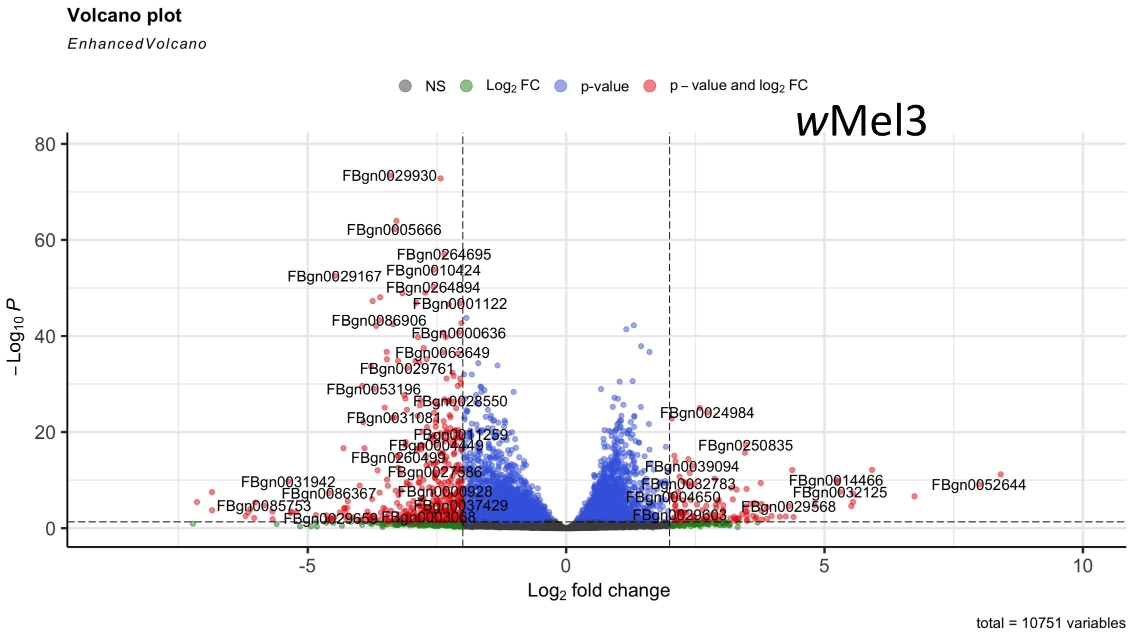


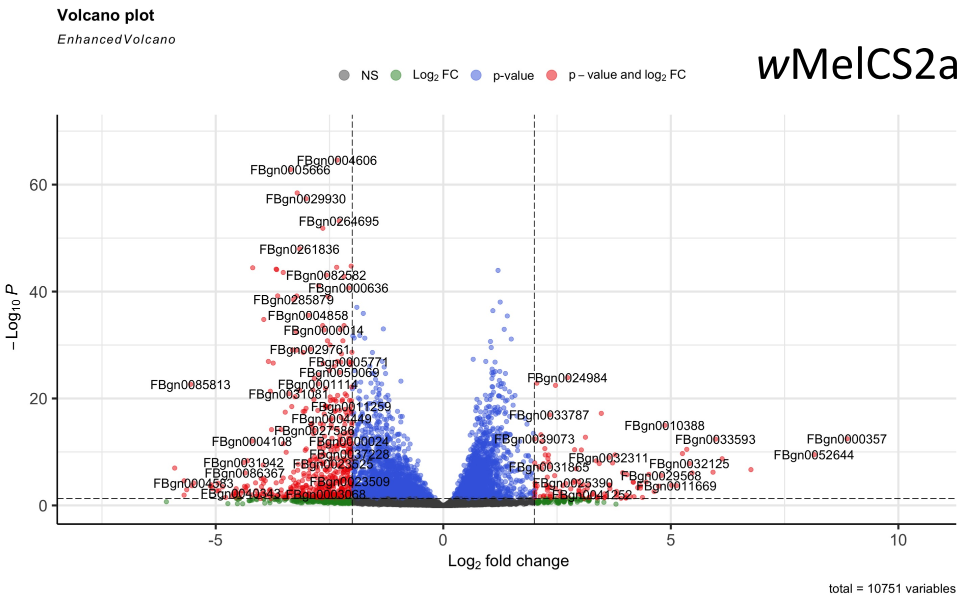


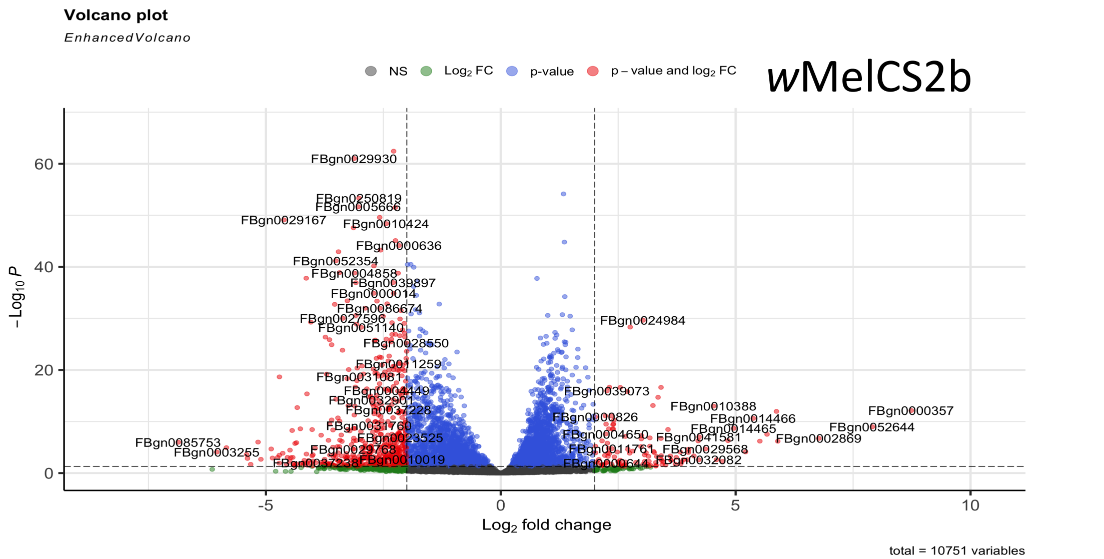
